# MAMBAxBrain: A Multi-task Neural Framework Linking Brain Functional Dynamics to Individual Fingerprints, Cognitive and Disease States

**DOI:** 10.64898/2026.02.08.704658

**Authors:** Yuqing Xia, Fahimeh Arab, Urmi Saha, Benjamin Sipes, Grayson Gooden, Minghan Chen, Ashish Raj

## Abstract

Functional magnetic resonance imaging (fMRI) contains rich individual, cognitive, and pathological information, yet no universal model exists for multi-task modeling of these dimensions. Here, we introduce MAMBAxBrain, a multi-task neural framework that integrates Mamba architecture with functional connectivity analysis to jointly model the temporal dynamics and spatial coordination of neural activity. MAMBAxBrain achieves high accuracy across four distinct fMRI objectives, including brain fingerprinting, cognitive task decoding, reaction time prediction, and schizophrenia classification, consistently outperforming state-of-the-art methods with robust cross-session generalization. Interpretability analyses show that each task engages distinct, biologically plausible circuitry: from higher-order association cortex for identity to subcortical–motor loops for reaction time and disrupted control-sensory connectivity for schizophrenia. These findings inform a longstanding debate: rather than operating through wholly separate or entirely shared systems, the brain preferentially recruits task-specific circuits while retaining common representational structure across functions.

## 1 Introduction

Functional magnetic resonance imaging (fMRI) has become a cornerstone of human neuroscience research, providing a noninvasive window into brain activity through blood oxygen level–dependent (BOLD) signals (Kwong et al., 1992). Fluctuations in these signals reflect the temporal evolution of neural dynamics within and between brain regions. By quantifying statistical dependencies among regional time series, researchers construct functional connectivity (FC) networks that characterize the brain’s large-scale communication patterns during rest and task performance (Biswal et al., 1995; Fox & Raichle, 2007; Friston, 2011; Greicius et al., 2003). Mounting evidence shows that fMRI contains rich information across multiple levels of brain organization, offering insights into neural function and behavior. Recent studies have further demonstrated that fMRI signals embed rich and decodable information about cognitive and perceptual processes. Models such as BrainGNN and MindBridge have shown that machine learning frameworks can effectively leverage these representations to decode brain states and even reconstruct presented stimuli with high accuracy (Li et al., 2021; S. Wang et al., 2024). In the spatial dimension, FC quantifies coordinated activity among brain regions and delineates intrinsic resting-state networks such as the default mode network, salience network, and central executive network (Fox & Raichle, 2007; Greicius et al., 2003). FC patterns also exhibit high specificity and reproducibility, serving as intrinsic “functional fingerprints” that facilitate accurate individual identification across scan sessions (Finn et al., 2015; Horien et al., 2019) and predict individual differences in cognitive behavior (Sieler et al., 2025; Yoo et al., 2018).

These findings suggest that BOLD-derived FC not only reflects the brain’s functional architecture but also carries individual-specific signatures, forming the basis of functional connectome fingerprinting (Finn et al., 2015). In the temporal dimension, BOLD time series capture neural dynamics. Temporal features such as the onset latency and duration (width) of the evoked hemodynamic response track the timing and persistence of underlying neural activity. These features significantly correlate with behavioral performance; specifically, delayed BOLD onsets and prolonged response durations are associated with slower trial-by-trial reaction times (Liu et al., 2004; Yarkoni et al., 2009). Clinical studies further reveal that disrupted FC organization accompanies psychiatric and neurological disorders (Steardo Jr et al., 2020), indicating that fMRI signals jointly encode stable individual identity, task-evoked cognitive states, and disease-specific dysconnectivity. Importantly, these dimensions rely on distinct spatiotemporal properties: unique, steady-state connectivity profiles define individual identity, whereas shared, temporally varying network reconfigurations support cognitive demands and characterize pathological disruptions (Calhoun et al., 2014; Cohen, 2018; Sanfratello et al., 2019). This calls for computational models that can jointly capture spatial connectivity and temporal dynamics within a joint spatiotemporal framework.

With the rise of deep learning, numerous models have been applied to fMRI to decode neural representations more efficiently. FC-based methods have achieved promising functional connectome fingerprinting (Amico & Goñi, 2018; Finn et al., 2015) and reliable cognitive-state decoding (Shirer et al., 2012). Recent models such as Brain Network Transformer (BrainNetTF) (Kan et al., 2022) and SPDNet (Huang & Van Gool, 2017) further improved FC representation from graph-structured and manifold learning perspectives. Yet these methods primarily rely on functional connectivity and overlook temporal dynamics of neural activity. Time-series-based approaches provide a complementary perspective by preserving activation timing, duration, and dynamic state transitions that are averaged out in static FC (Allen et al., 2014). This ability to leverage the full sequence has led to strong decoding performance across diverse tasks, including cognitive states (X. Wang et al., 2020) and individual identification (Sarar et al., 2021), and clinical diagnosis of schizophrenia (Yan et al., 2019). Nevertheless, these temporal-only models often ignore FC structure, which reduces interpretability and makes it difficult to link the learned representations to known functional networks or connectivity-based biomarkers.

Recent Mamba-based architectures extend this temporal modeling capability to fMRI. Mamba is a selective state-space model (SSM) that achieves linear-time sequence modeling while preserving the ability to learn long-range dependencies. It can scale to very long sequences with stable gradients. These properties make Mamba particularly suitable for fMRI, where BOLD signals form long multi-regional time series and require modeling both rapid fluctuations and delayed hemodynamic responses. Building on this foundation, several recent fMRI-specific Mamba variants have been proposed. Brain-Mamba (Behrouz & Hashemi, 2024) models voxelwise BOLD fluctuations using selective state-space dynamics, and Causal fMRI-Mamba (Deng et al., 2025) targets task-state decoding via causal temporal representations. FST-Mamba (Wei et al., 2025) further integrates spatial topology and temporal dynamics through a hierarchical state-space formulation. Nevertheless, its design is tailored to specific decoding objectives and does not generalize effectively across diverse analytical tasks.

Overall, most existing deep learning approaches still address either functional connectivity or temporal dynamics in isolation, which leaves the full spatiotemporal structure of fMRI underutilized. In addition, many models are designed for a single task with limited scalability, and the absence of a general spatiotemporal learning framework restricts the transfer of learned features to new settings. This gap highlights the need for a scalable framework capable of learning generalizable neural representations from both temporal dynamics and functional connectivity and adapting efficiently to new tasks via few-shot adaptation. Such a framework would not only reduce the need to train separate models for each new study, but also enable more consistent biomarkers that link individual identity, cognitive state, behavior, and pathology within a common representational space.

To address the above challenges, we propose MAMBAxBrain, a multi-task neural framework that integrates temporal and connectivity modeling within a dual-stream architecture. Designed for low computational complexity and efficient handling of long BOLD sequences, MAMBAxBrain generalizes readily across sessions and supports few-shot adaptation to new tasks. We evaluate the framework across a broad set of fMRI-based analyses and further assess its cross-session generalizability and multi-task performance within a unified backbone. A schematic of the proposed model, data streams, and input/output specifications is shown in Fig. 1.

**Figure 1:**
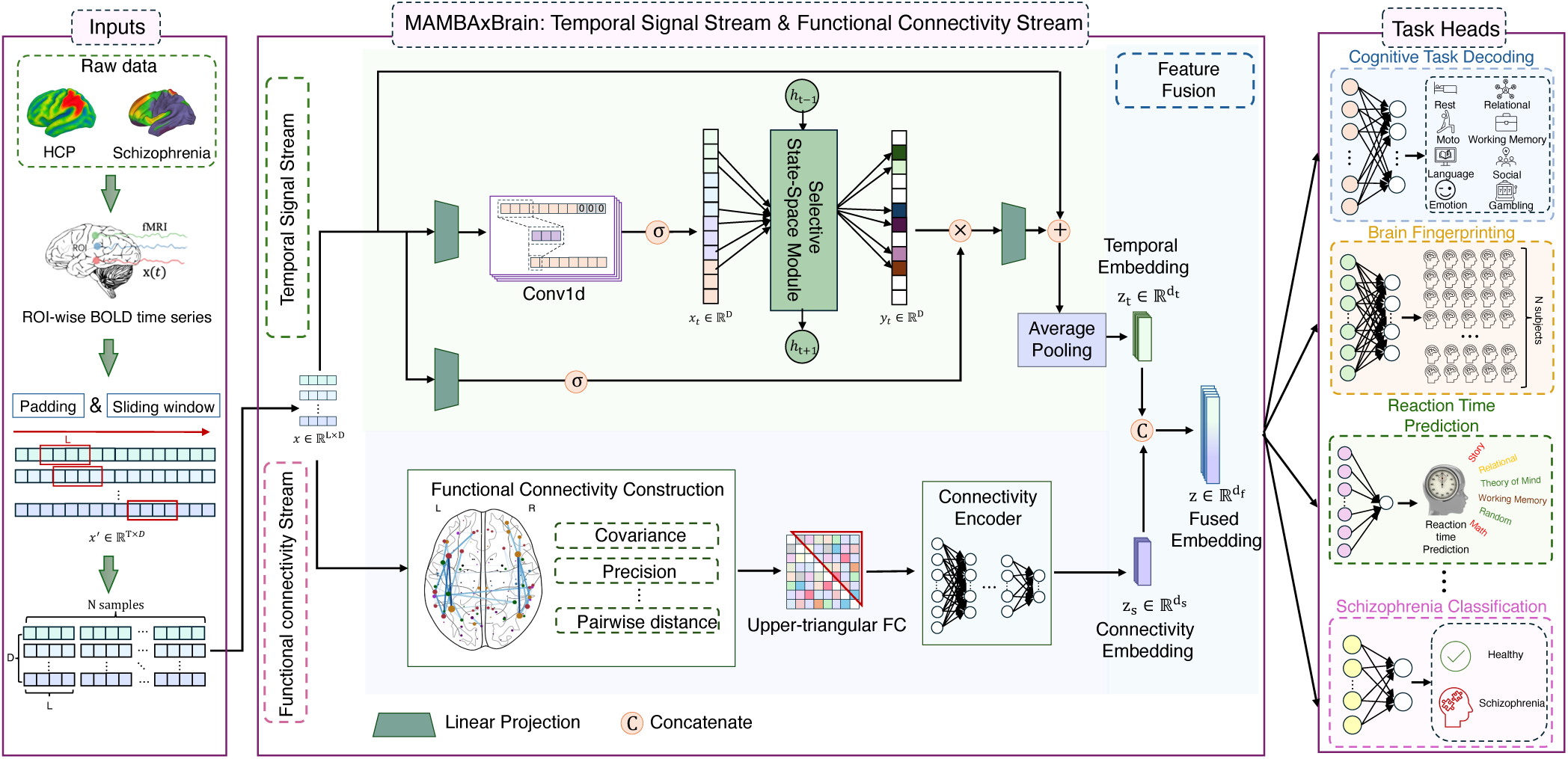
Overview of our proposed MAMBAxBrain framework. Input: ROI-wise BOLD time series are segmented into multiple samples using a sliding-window strategy, followed by zero padding for length alignment and z-score normalization across regions. Dual-stream core: a temporal signal stream based on a selective SSM that captures long-range temporal dependencies, and an FC stream that encodes inter-regional relationships from pairwise connectivity matrices. The embeddings from both streams are fused into a unified representation. Task Heads: The shared representation feeds multiple lightweight task heads that perform brain fingerprinting, cognitive task decoding, reaction time prediction, and schizophrenia classification, enabling unified and scalable multi-task fMRI modeling.

Our main contributions are as follows:

1. **A dual-stream spatiotemporal framework:** We propose a dual-stream neural architecture that integrates temporal dynamics and functional connectivity into a shared representational space. The Mamba-based temporal stream captures long-range dependencies in BOLD signals, while the functional connectivity stream encodes interregional coactivation patterns. A fusion module combines these complementary features into joint spatiotemporal representations that support diverse downstream tasks. Unlike prior fused models such as FST-Mamba, which are designed for single-task learning using multiple state-space encoders over dynamic connectivity sequences, MAMBAxBrain provides a streamlined, unified backbone with a learnable fusion module that enables multi-task learning and few-shot transfer across tasks.
2. **Superior multi-task performance and robust generalization.** MAMBAxBrain supports multi-task learning within a unified backbone and consistently outperforms state-of-the-art benchmark methods in four representative tasks: brain fingerprinting, cognitive task decoding, reaction time prediction, and schizophrenia classification. Cross-session and multi-task evaluations further demonstrate its robustness to inter-session variability and its ability to generalize across new tasks.
3. **Interpretability and neurological insights:** Model interpretation reveals shared yet dissociable network signatures. We provide a mechanistic understanding by showing that individual identity is encoded in higher-order control, default mode, and dorsal attention networks, cognitive tasks are decoded from sensory-motor systems, reaction time is predicted by a subcortical-motor-limbic circuit, and distinguishes schizophrenia through disruptions in top-down control circuits linking frontoparietal and cingulo-opercular hubs to auditory-visual streams.

## 2 Results

### 2.1 Experimental overview

Details of fMRI data acquisition and preprocessing are provided at the beginning of the Methods section, and implementation details, dataset configurations, and optimization settings are described in the Experimental Settings subsection. To evaluate the single-task performance, cross-session generalization, and multi-task generalization of MAMBAxBrain, we conducted three types of experiments spanning four representative tasks: brain fingerprinting (FP), which identifies unique individuals; cognitive task decoding (TD), involving the classification of seven distinct cognitive tasks including motor, working memory, language, gambling, social, relational, emotion tasks and a resting-state; reaction time prediction (RT), a regression task estimating condition-level mean reaction time; and schizophrenia classification (SZ), a binary classification distinguishing patients from healthy controls. We report three types of experiments on our study cohort: (1) Single-task experiments to evaluate model performance within each task on the Human Connectome Project Young Adult (HCP-YA) dataset, referred to as in-domain data. (2) Cross-session experiments to assess generalization by training on HCP-YA dataset and testing on independent session data. For resting-state fMRI, the first two HCP-YA runs were used for in-domain training, and the remaining two runs were used for cross-session evaluation. For all other states of fMRI (Working Memory, Relational, etc.), cross-session generalization was assessed using the Human Connectome Project Test-Retest (HCP-RT) dataset, collected from 45 participants during a second full fMRI session approximately 4–6 months later. (3) Multi-task experiments to examine the framework’s performance and few-shot transfer to held-out tasks. Due to differences in atlas parcellation, the schizophrenia dataset was excluded from the multi-task experiments.

### 2.2 Single-task learning and cross-session generalization

To comprehensively evaluate the proposed framework, we compared MAMBAxBrain with several representative benchmarking methods across four independent tasks, including SPDNet (Huang & Van Gool, 2017), Brain Network Transformer (BrainNetTF) (Kan et al., 2022), CorrNN (Sarar et al., 2021), Mamba (Gu & Dao, 2024), and FST-Mamba (Wei et al., 2025). These methods span three typical input paradigms: FC-only models that use FC as input (SPDNet, BrainNetTF, CorrNN), time series-only models that use time series as input (Mamba), and models that jointly use time series and FC (Both) (FST-Mamba and MAMBAxBrain). Table 1 summarizes the quantitative results for in-domain evaluation on the HCP-YA dataset and cross-session generalization on the HCP-RT dataset, while Fig. 2 visualizes the overall performance trends. A detailed discussion of the input features, architectural differences, and limitations of each baseline is provided in Section Analysis of baseline models.

**Figure 2:**
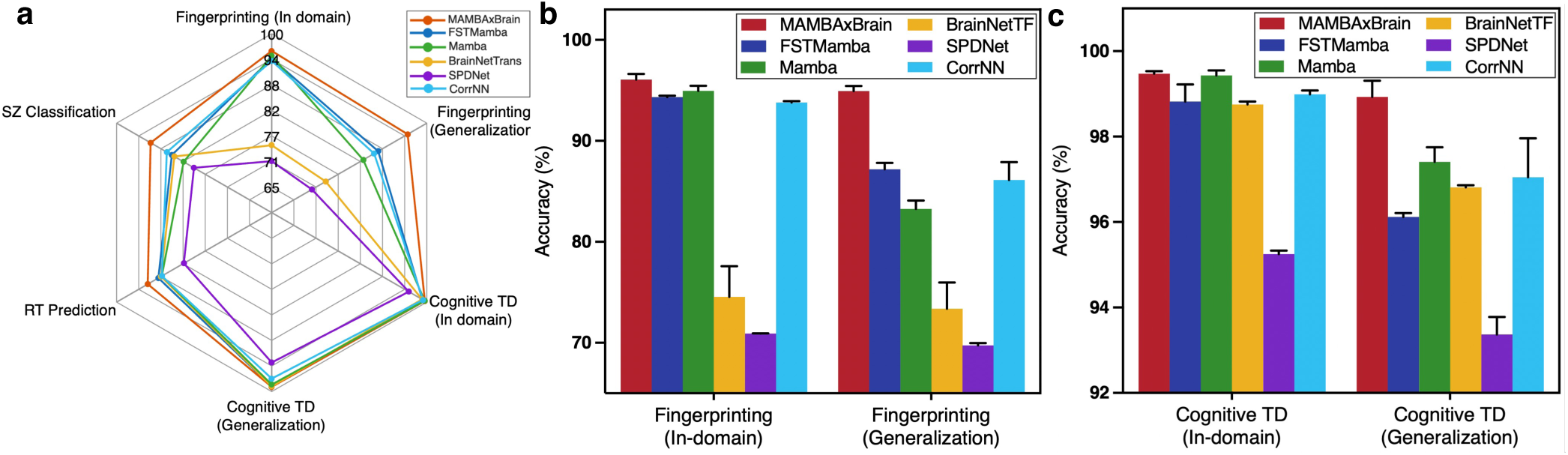
Overall single-task performance and cross-session generalization across models. **a)** Spider chart comparing performance of SPDNet, BrainNetTF, Mamba, FST-Mamba, and MAMBAxBrain. **b, c)** Benchmarking performance on brain fingerprinting and cognitive task decoding (TD) under in-domain evaluation on HCP-YA and cross-session generalization on HCP-RT.

**Table 1:**
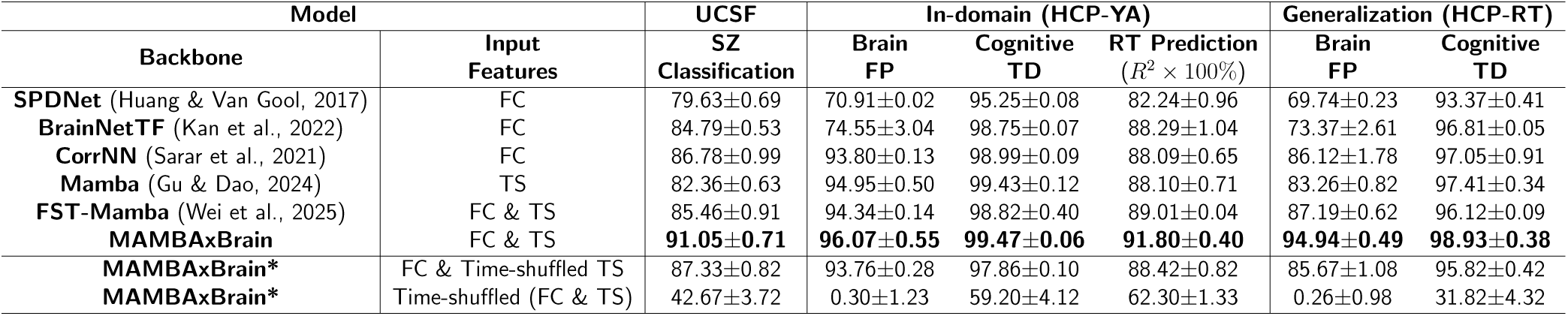
Comparison of individual task performance across methods for in-domain evaluation and cross-session generalization. Accuracy is reported for classification tasks: brain fingerprinting (FP), cognitive task decoding (TD), and schizophrenia classification (SZ), while the *R*^2^ score is reported for regression task: reaction time prediction (RT). TS denotes time series. FC denotes functional connectivity. Time-shuffled controls: global time-series shuffle (FC preserved) vs. ROI-wise shuffle (FC recomputed).

Overall, MAMBAxBrain achieves the best performance across all tasks and evaluation settings, with particularly pronounced advantages in cross-session generalization. In the in-domain (HCP-YA) evaluation, it reaches 96% accuracy on brain fingerprinting, outperforming both FC-only models (e.g., SPDNet, BrainNetTF) and the TS-only model (Mamba). For cognitive task decoding, it attains 99% accuracy with reduced variance, indicating more stable training and stronger reproducibility even when overall accuracy is already high. For reaction time prediction, it achieves an *R*^2^ of 0.92, demonstrating its effectiveness in capturing complex dynamics associated with behavior. For schizophrenia classification, the model reaches 91% accuracy, clearly outperforming FC-only, TS-only, and existing FC-TS fusion approaches. Notably, FC-only methods can achieve relatively strong performance on some tasks (e.g., cognitive task decoding), but their overall disadvantages become more evident in individual identification and cross-session settings, suggesting that relying solely on static connectivity patterns is insufficient to capture subject-specific dynamic differences and robust cross-session representations. In contrast, the TS-only Mamba performs strongly in-domain but exhibits a larger drop under cross-session (HCP-RT) evaluation, indicating that modeling time-series dynamics alone can remain unstable under session shifts. By comparison, methods that leverage both TS and FC are generally more robust, and MAMBAxBrain further improves performance through a more effective and unified FC–TS integration. In contrast, FST-Mamba adopts a complex hierarchical design with two Mamba encoders and multiple sequential aggregation operations, resulting in substantially higher computational overhead and longer training time, which may hinder optimization and reduce robustness relative to our streamlined architecture. This efficiency difference is quantified in Supplementary Table S5: under matched inputs MAMBAxBrain uses roughly half the FLOPs (10.9 vs. 28.0 G) and training time per step (26.3 vs. 60.0 ms) of FST-Mamba, while remaining close to the single-stream Mamba backbone (10.9 vs. 10.6 G FLOPs).

In the cross-session (HCP-RT) evaluation, the advantages of MAMBAxBrain become even more pronounced: it achieves 94.94% accuracy on brain fingerprinting, improving by approximately 9 percentage points over the best-performing baseline. For cognitive task decoding, it maintains a near-ceiling accuracy of 98.93% under cross-session conditions and remains clearly superior to competing models. These results suggest that jointly leveraging the temporal dynamics in time series and the cross-region coordination captured by FC facilitates learning more stable and transferable representations, thereby improving cross-session generalization.

To further validate the critical role of temporal structure in time series, we introduce two time-shuffled settings as temporal-disruption controls (Table 1). In the global time-series shuffle setting, a single random permutation is applied to the temporal order of the input time series and shared across all ROIs, while the FC input is kept unchanged. Under this condition, performance consistently degrades across tasks and evaluation settings; the most noticeable drops are observed for schizophrenia classification on UCSF, decreasing from 91.05% to 87.33%, and for cross-session brain fingerprinting on HCP-RT, decreasing from 94.94% to 85.67%. Because static FC is preserved, this ablation isolates the contribution of temporal order in the time-series stream, indicating that temporal ordering provides additional discriminative information to MAMBAxBrain. In the ROI-wise shuffle setting, the temporal order within each ROI is independently permuted, disrupting cross-ROI temporal alignment and altering static FC. As both TS and FC inputs are perturbed, this results in a dramatic performance collapse across all tasks.

We also assessed the stability of these results with respect to the sliding-window evaluation. Under a stride of 1, adjacent windows overlap by 99 of their 100 TRs; repeating the fingerprinting and cognitive-decoding evaluations with non-overlapping windows (stride equal to the window length) changed accuracy by less than one percentage point in every setting, both in-domain and across sessions (Table S2). Together with the consistency observed across folds, this indicates that performance reflects stable temporal signatures and is not driven by the degree of overlap between adjacent windows.

To establish the contribution of each architectural component, we further performed a controlled, change-one-thing-at-a-time ablation in which every variant keeps the rest of the architecture and the training budget fixed (Supplementary Table S8). Using only the temporal stream (equivalent to the Mamba backbone) or only the FC stream (equivalent to CorrNN) reduced performance, most markedly for cross-session fingerprinting (94.94 to 83.26 and 86.12, respectively), confirming that the two streams are complementary. Replacing the learnable fusion with simple concatenation produced a smaller but consistent decline, and substituting the Mamba temporal encoder with Transformer, TCN, LSTM, or GRU encoders of matched capacity degraded performance monotonically, again with the largest gaps under cross-session evaluation. These results indicate that the gains arise jointly from the selective state-space encoder, the dual-stream design, and the learnable fusion, rather than from any single factor or from model capacity alone.

### 2.3 Multi-task learning

We further evaluated its ability to support multi-task learning and generalization. Our model was trained on base tasks and then few-shot adapted to held-out tasks by fine-tuning on 20% of the target-task data (few-shot transfer, not zero-shot generalization) to examine how effectively the learned representations adapted to new tasks. As summarized in Fig. 3a, brain fingerprinting served as the base objective, and additional objectives (cognitive task decoding and reaction time prediction) were sequentially added to evaluate performance on the HCP-YA dataset under multi-task configurations. When trained solely on brain fingerprinting, the model achieved 96% accuracy on that task. However, its performance dropped substantially when directly applied to held-out tasks without any fine-tuning (zero-shot) such as cognitive task decoding and reaction time prediction, indicating that single task representations were more specialized and lacked generality. Joint training on fingerprinting and cognitive decoding improved performance on both tasks (95% and 99%, respectively) and markedly enhanced the reaction time prediction to *R*^2^ = 0.72. This suggests that incorporating cognitive decoding helps the model to capture task-relevant temporal dynamics while preserving individual-specific features, thereby improving cross-task transfer. Crucially, fingerprinting and cognitive decoding are trained on the same input data, namely the identical HCP-YA scans and sliding windows, and differ only in their labels, subject identity versus cognitive task. The increase in reaction-time prediction when task decoding is added to training (RT *R*^2^ rising from 0.24 to 0.72; Fig. 3b) therefore cannot be attributed to additional training data, since the data are identical; it instead reflects the auxiliary task-decoding objective shaping more transferable representations. When reaction time prediction was further included during training, the model maintained high accuracy across all tasks and further improved reaction time prediction to *R*^2^ of 0.91. These results demonstrate that integrating complementary tasks promotes the learning of richer, more transferable spatiotemporal representations.

**Figure 3:**
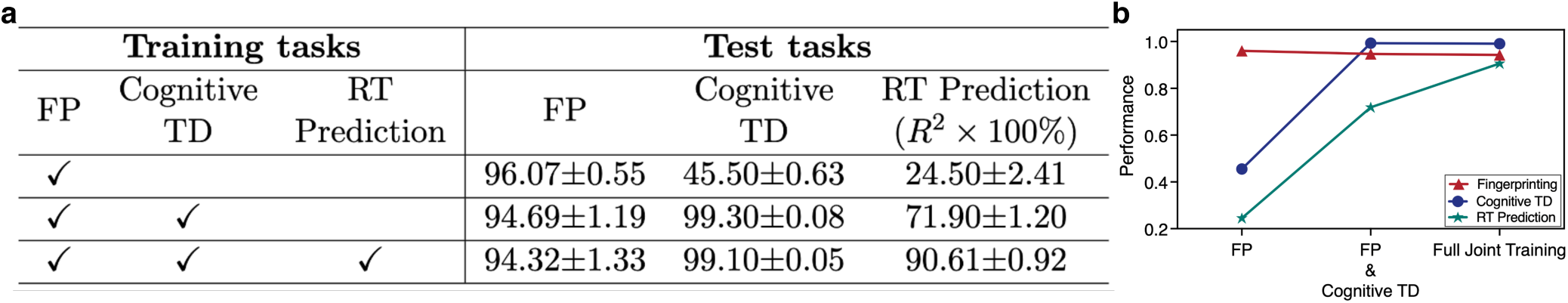
Evaluation of multi-task performance and few-shot adaptation on the HCP-YA Dataset. Here, FP denotes Fingerprinting, TD denotes Task Decoding, and RT denotes Reaction Time. **a)** Performance of models trained under multi-task configurations. The symbol ✓ indicates tasks included during training, while blank entries represent few-shot adaptation to held-out tasks (20% of the target task is used for fine-tuning, i.e., few-shot transfer rather than zero-shot generalization). **b)** Performance trends with increasing numbers of training tasks.

Because every baseline in Table 1 is trained single-task, we further separated MAMBAxBrain’s advantage into an architectural and a multi-task contribution at a matched budget: every method was retrained single-task from scratch on the same 20% of each target task, and the multi-task MAMBAxBrain was additionally few-shot adapted on that same 20%. As shown in Supplementary Table S4, among the single-task models MAMBAxBrain is the best on every task (e.g., RT *R*^2^ × 100 = 34.50 versus ≤ 31.20 for all baselines), isolating the architectural advantage at the few-shot budget; and few-shot adapting the multi-task model on the same 20% adds a further, larger gain (e.g., RT 34.50 → 71.90, FP 70.92 → 84.20), which isolates the multi-task benefit because the architecture and target data are held fixed. The improvement over baselines therefore reflects both a stronger architecture and genuine cross-task transfer rather than data volume alone.

Fig. 3b illustrates performance trends with increasing number of tasks. Overall, as training tasks increases, MAMBAxBrain performance remained stable or even improved consistently across tasks, demonstrating strong multi-task learning and scalability. To further interpret the model’s scalable behavior, we examined the functional connectivity patterns learned under joint multi-task training across all tasks (Supplementary Fig. S2). The resulting connectivity maps revealed a comprehensive, integrated network organization. This multi-task model learned a composite set of features, simultaneously leveraging the higher-order (control, default mode, and dorsal attention), sensory-motor, and subcortical-limbic connections required to optimize performance across all tasks.

### 2.4 Task-specific performance and analysis

#### 2.4.1 Brain fingerprinting

We first assessed MAMBAxBrain’s capacity to capture stable, individual-specific neural signatures using the brain fingerprinting task. Fig. 4a shows its classification accuracies across eight cognitive tasks under in-domain evaluation (HCP-YA) and cross-session generalization (HCP-RT). In the in-domain setting, our model achieved high accuracy with low variance across tasks, indicating stable discriminative ability under diverse cognitive paradigms. Under cross-session evaluation, average accuracy decreased slightly and variability increased, reflecting distributional shifts between sessions. Most tasks, such as working memory (WM), language, gambling, and emotion, retained relatively high accuracy, suggesting that the model captures stable and transferable functional connectivity patterns associated with these cognitive processes. By contrast, tasks such as social and relational showed more noticeable performance declines, implying that their associated activation patterns may vary more across scanning sessions. Overall, our model exhibits minimal performance variation across tasks and maintains a stable overall trend, demonstrating strong temporal generalization capability across multiple cognitive conditions.

**Figure 4:**
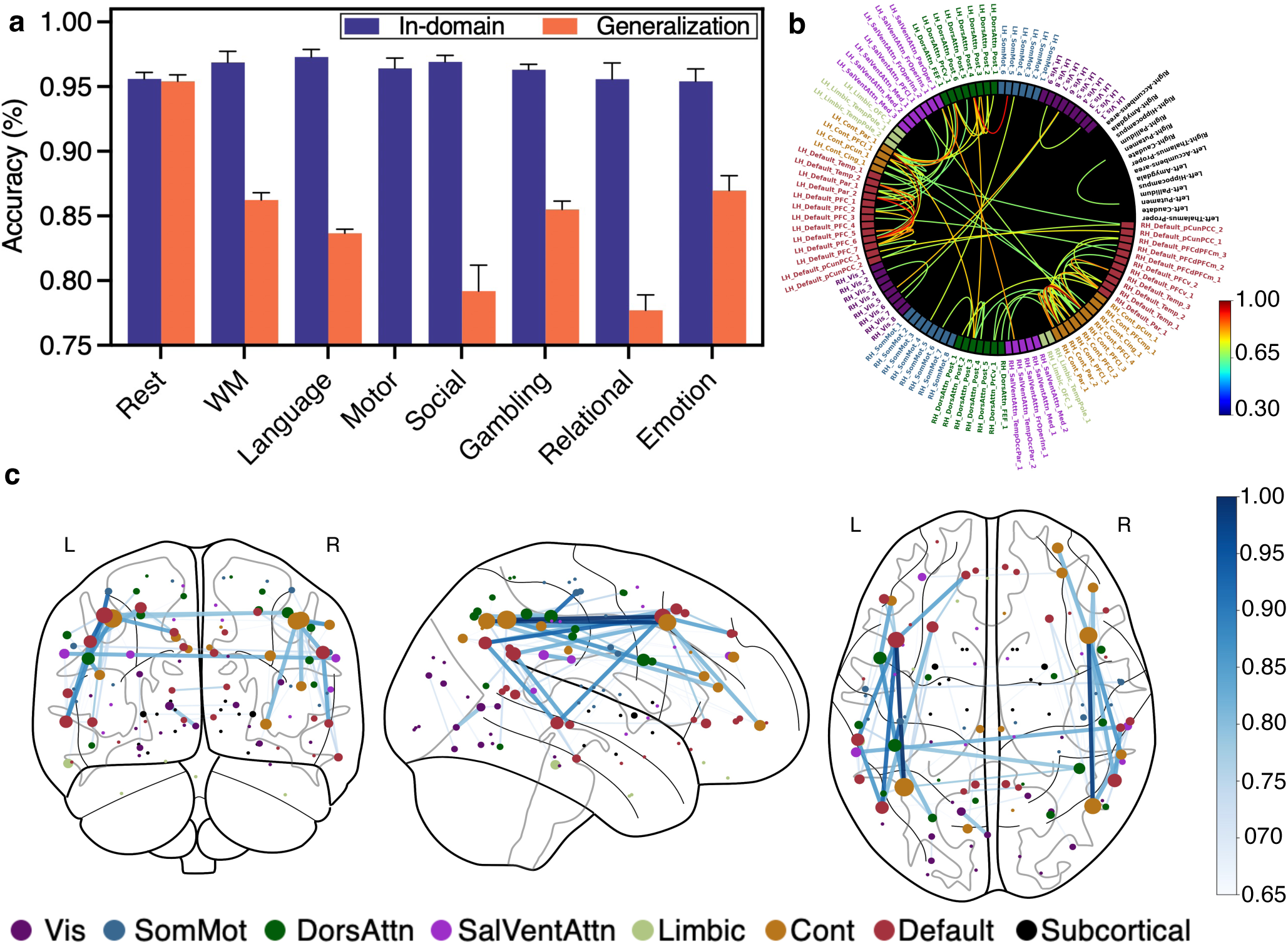
Brain fingerprinting performance and learned functional connectivity patterns. **a)** Individual identification accuracy across cognitive tasks under in-domain evaluation (HCP-YA) and cross-session generalization (HCP-RT). The HCP-YA evaluation includes approximately 761 participants per task on average. For HCP-RT, the Rest task includes 732 participants, whereas all other tasks contain 45 participants. The motor task is not available in HCP-RT and is therefore omitted. **b)** Connectivity circle showing functional connections ranked by edge importance (threshold = 0.65), with parcel labels colored by their functional network membership. **c)** Connectome map projected onto atlas coordinates, showing the spatial distribution of high-importance functional connections (threshold on edge importance = 0.65). Edge thickness is proportional to the learned connection importance score, and node size reflects node strength computed by summing the importance weights of adjacent edges in the thresholded importance graph.

To identify the functional connectivity features contributing to individual identification, edge-importance weights were derived from MAMBAxBrain’s learned functional connectivity representations. The methodological details of feature importance estimation and back-projection are provided in Section Feature importance analysis. The results reveal that the model’s predictive strategy is concentrated in higher-order association cortices. Specifically, the connections identified as most critical for fingerprinting were those within and between the default mode, control, and dorsal attention networks. These findings are visualized in Fig. 4b,c. The connectivity circle (Fig. 4b) highlights the dominance of the connectivities involving the control, default mode and dorsal attention networks. In this plot, brain region labels are colored by their functional network membership. The connectome map (Fig. 4c) projects these same importance scores onto atlas coordinates, confirming their spatial concentration in these higher-order systems. In Fig. 4c, the edge thickness represents the learned importance of each connection, while the node size reflects the node strength in the importance-weighted connectome, where edges are weighted by their importance scores rather than raw connectivity. Specifically, we first threshold the connection importance scores and retain only edges above the threshold, then compute each node’s total importance by summing the importance weights of all retained edges incident to that node. These interpretability results demonstrate that MAMBAxBrain learns a sparse feature set that reflects interpretable connectivity patterns. These MAMBAxBrain identified networks align with prior findings (Finn et al., 2015), suggesting that our model inductively derives these networks as sources of individual-specific connectome “fingerprints”.

#### 2.4.2 Cognitive task decoding

We next investigated how MAMBAxBrain distinguishes cognitive states across tasks. The influence of temporal granularity on classification performance was first examined by varying the sliding-window parameters. Fig. 5a shows the effect of window size (i.e., the number of consecutive fMRI time points used per sample) on classification accuracy, evaluated on both in-domain and generalization datasets. Accuracy increases with window length up to 100 time points before declining, suggesting that excessively large windows introduce redundant or cross-task information, while overly short windows fail to capture sufficient temporal structure. Thus, a moderate window size balances temporal detail and statistical stability, yielding optimal performance. Complementary to window size, the sliding stride determines the temporal overlap between consecutive samples. As shown in Fig. 5b, increasing stride length leads to a gradual decline in classification accuracy, particularly in cross-session evaluations. Larger strides reduce temporal sampling density, causing the model to miss fine-grained neural dynamics and weakening generalization. Smaller strides, by contrast, provide denser temporal coverage and smoother transitions between windows, enhancing stability and robustness. We next examined task-level discrimination using a confusion matrix and t-SNE visualization. Fig. 5c shows that all eight cognitive tasks achieve high accuracy (*>*99%), with few misclassifications. Fig. 5d visualizes task embeddings in a t-SNE space, revealing distinct clusters for eight tasks. Minor overlaps between gambling and relational tasks suggest shared activation patterns, reflecting similarities in cognitive control and reward processing.

**Figure 5:**
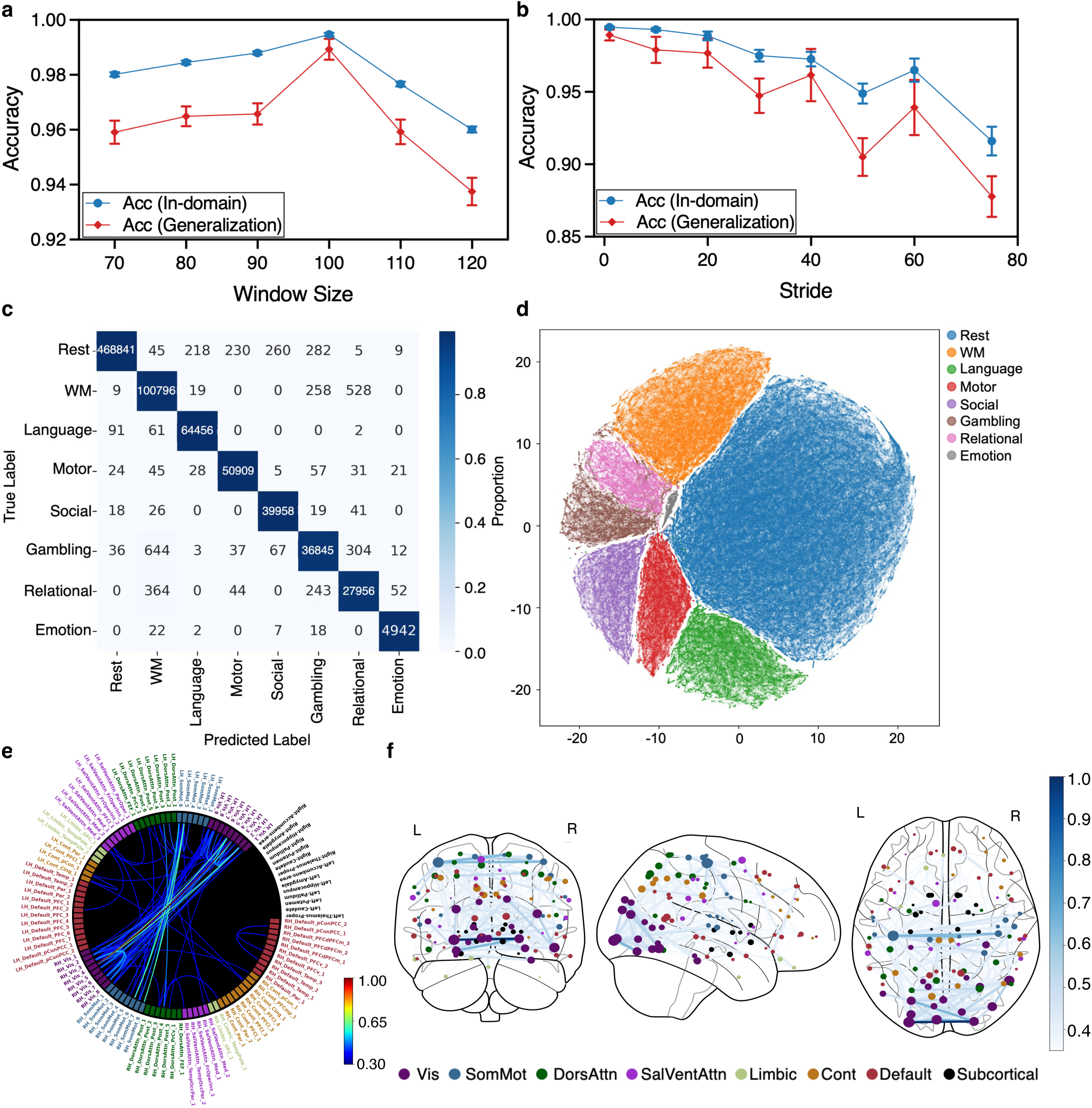
Cognitive task decoding performance and learned functional connectivity patterns. **a, b)** Effect of window size and stride on in-domain (HCP-YA) and generalization (HCP-RT) performance. **c)** Confusion matrix of cognitive task classification. **d)** t-SNE plot of learned task representations. **e)** Connectivity circle showing functional connections ranked by edge importance (threshold = 0.35), with parcel labels colored by their functional network membership. **f)** Connectome map projected onto atlas coordinates, depicting the spatial distribution of highly weighted connections (threshold = 0.35). Edge thickness is proportional to the learned connection importance score, and node size reflects each region’s node strength computed over the thresholded importance graph.

Analysis of edge-importance weights uncovered network patterns distinguishing different cognitive states. In stark contrast to brain fingerprinting, the model relied primarily on sensory, motor, and attention networks, reflecting their dominant role in task-specific processing. Specifically, the connections identified as most critical for task decoding were those within and between the visual, somatomotor, and dorsal attention networks. These findings are visualized in Fig. 5e,f. The connectivity circle (Fig. 5e) highlights the dominance of these sensory-motor and attention systems, with parcel labels colored by their functional network membership. The connectome map (Fig. 5f) projects these same importance scores onto atlas coordinates, confirming their spatial concentration in these task-positive regions. As in the previous analysis, the edge thickness represents the importance of each specific connection, while the node size reflects the total importance of that region (i.e., its nodal degree in the importance map). This clear dissociation, also summarized in the network-level analysis (Fig. 8b), demonstrates that the model correctly identifies that “who” a person is and “what” they are doing are encoded in distinct neural-functional substrates. It is also intriguing that identity-governing FCs are dominantly ipsilateral while task-governing FCs are strongly inter-hemispheric.

#### 2.4.3 Reaction time prediction

We further tested the model’s ability to predict behavioral variability in reaction time. In Fig. 6a, predicted reaction times align closely with the diagonal, indicating strong correspondence with ground-truth values (*R*^2^ = 0.92). A few noticeable outliers appear primarily in the random (social) and relational tasks. To further assess prediction agreement and potential systematic bias, we used a Bland–Altman analysis (Fig. 6b). The 95% limits of agreement (−697.4, 659.9 ms) fall within approximately ±25% of the average reaction time across participants, indicating acceptable agreement between predicted and observed data. The mean bias (−18.7 ms) is close to zero, suggesting no consistent overestimation or underestimation by the model. Task-specific differences were also observed: predictions for the working memory task show tightly clustered distributions, whereas other tasks exhibit broader dispersion and a few outliers, suggesting that the current model may less effectively capture the neural–behavioral coupling underlying these specific behaviors, potentially due to greater unmodeled behavioral variability.

**Figure 6:**
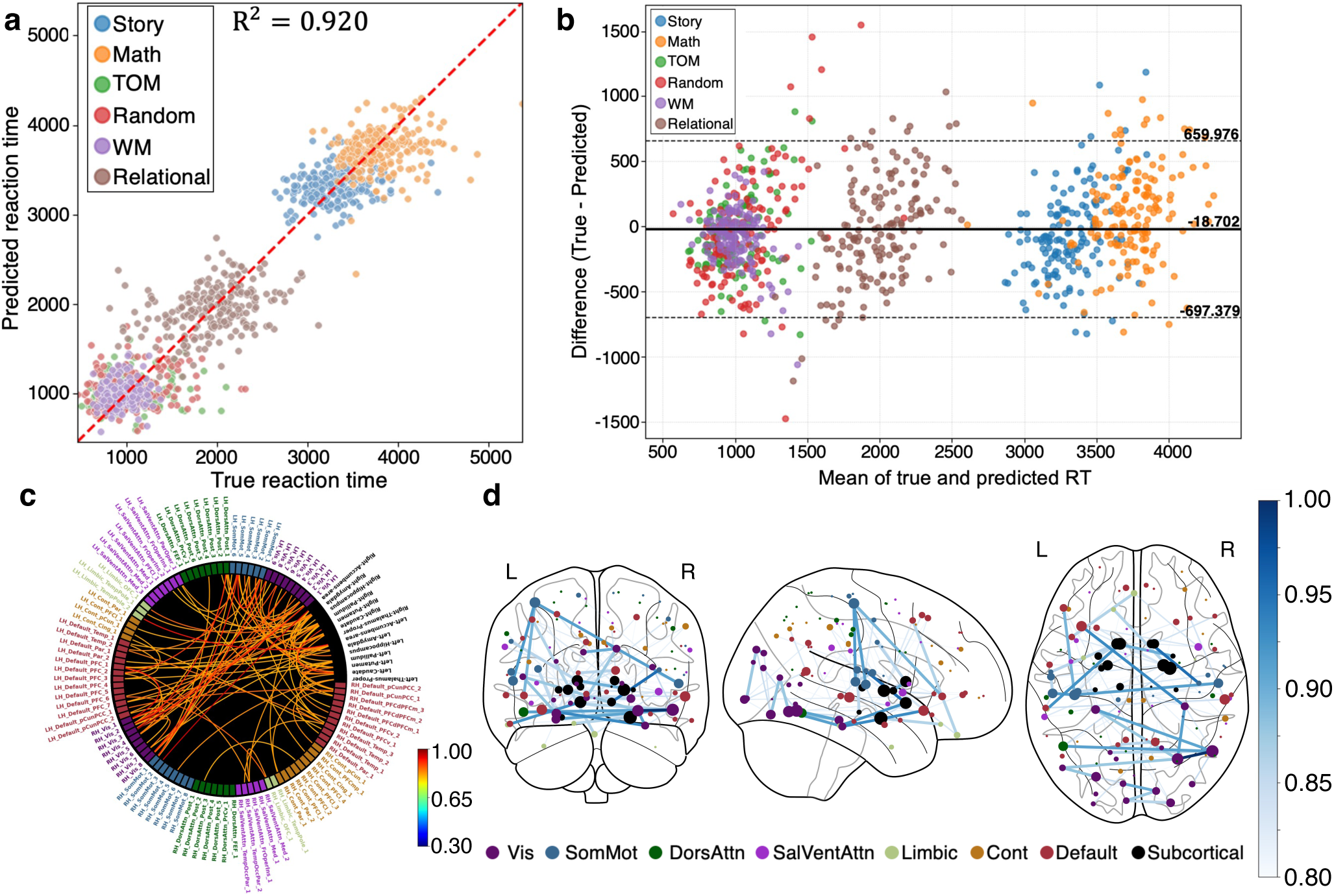
Reaction time prediction performance and learned connectivity patterns. **a)** Task-wise regression scatterplot, showing predicted versus true reaction times. **b)** Bland–Altman plot with mean reaction time on the x-axis and prediction error (True - Predicted) on the y-axis. **c)** Connectivity circle showing functional connections ranked by edge importance (threshold = 0.8). **d)** Connectome plot projected onto atlas coordinates, showing the spatial distribution of functional connections (threshold = 0.8). Edge thickness is proportional to the learned connection importance score, and node size reflects each region’s node strength computed over the thresholded importance graph.

Because the prediction target is the condition-level mean reaction time, this aggregate *R*^2^ = 0.92 is partly driven by systematic differences in mean RT between conditions rather than by individual neural variability alone. To isolate the within-condition signal, we additionally computed *R*^2^ separately within each condition (Supplementary Table S3): the model then explains substantially less variance (mean within-condition *R*^2^ ≈ 0.19, range 0.12–0.27). The strong aggregate fit should therefore be interpreted as primarily reflecting coarse between-condition structure, with individual within-condition reaction-time prediction remaining limited (see also Limitations).

Edge-importance mapping revealed a unique predictive circuit, distinct from both fingerprinting and task decoding. As shown in Fig. 6c,d, the model’s predictions were not driven by higher-order cortical networks, but by a specific “processing speed” circuit involving subcortical regions. The most critical features were connections from the subcortical network to the somatomotor, ventral attention, and limbic networks, as well as connections within the visual, limbic, and somatomotor networks themselves. These findings are visualized in the connectivity circle (Fig. 6c), which highlights the dominance of this subcortical-motor-limbic circuit, with parcel labels colored by their functional network membership. The connectome map (Fig. 6d) projects these same importance scores onto atlas coordinates, confirming the spatial distribution of these key features. This finding is also clearly reflected in the network-level analysis (Fig. 8c). This demonstrates the model’s ability to move beyond coarse state-classification and identify a fine-grained, biologically plausible circuit for predicting a specific behavioral outcome.

#### 2.4.4 Schizophrenia classification

Finally, to assess the model’s clinical applicability, we applied MAMBAxBrain to schizophrenia classification and evaluated the separability of learned representations between patients and healthy controls. In Fig. 7a, the t-SNE projection of the latent feature space reveals distinct clusters for the two groups, with partial overlap reflecting inter-individual heterogeneity in neural representations. We next assessed classification performance and potential group-level biases using the confusion matrix (Fig. 7b). Our model correctly identified 89 of 93 healthy controls and 40 of 45 patients, demonstrating strong discriminative performance with few misclassifications. Figure 7c shows the receiver operating characteristic (ROC) curve, with an area under the curve (AUC) of 0.95, confirming excellent separability between patients and healthy controls. Because the UCSF cohort is modest in size (45 patients, 93 controls), we further assessed the robustness of this result and directly addressed the possibility of within-subject leakage using diagnosis-stratified, subject-disjoint 5-fold cross-validation, in which no subject’s windows are shared between training and test. Under this stricter protocol MAMBAxBrain attains an accuracy of 82.14% ± 11.03% and an AUC of 0.83 ± 0.17 (mean ± s.d. across folds), the best among all models evaluated under the same protocol (Supplementary Table S6; strongest baseline FST-Mamba, 0.76/0.80). As anticipated by the reviewers, performance is lower than the within-split estimate (Table 1) and the fold-to-fold variability is large, reflecting the limited sample size; the schizophrenia results should therefore be regarded as promising but preliminary. To further test whether the performance is dataset-specific, we additionally evaluated the trained model, without retraining, on the independent COBRE cohort (72 patients, 74 controls; Supplementary Table S7), where MAMBAxBrain attained the best external performance among all models (AUC = 0.81, accuracy = 0.68) while showing the expected drop under cross-site domain shift, most notably in sensitivity (0.43). Together these analyses indicate that the learned schizophrenia-related representations generalize above chance beyond a single split and beyond a single cohort, although larger multi-site samples will be needed to establish robust clinical performance.

**Figure 7:**
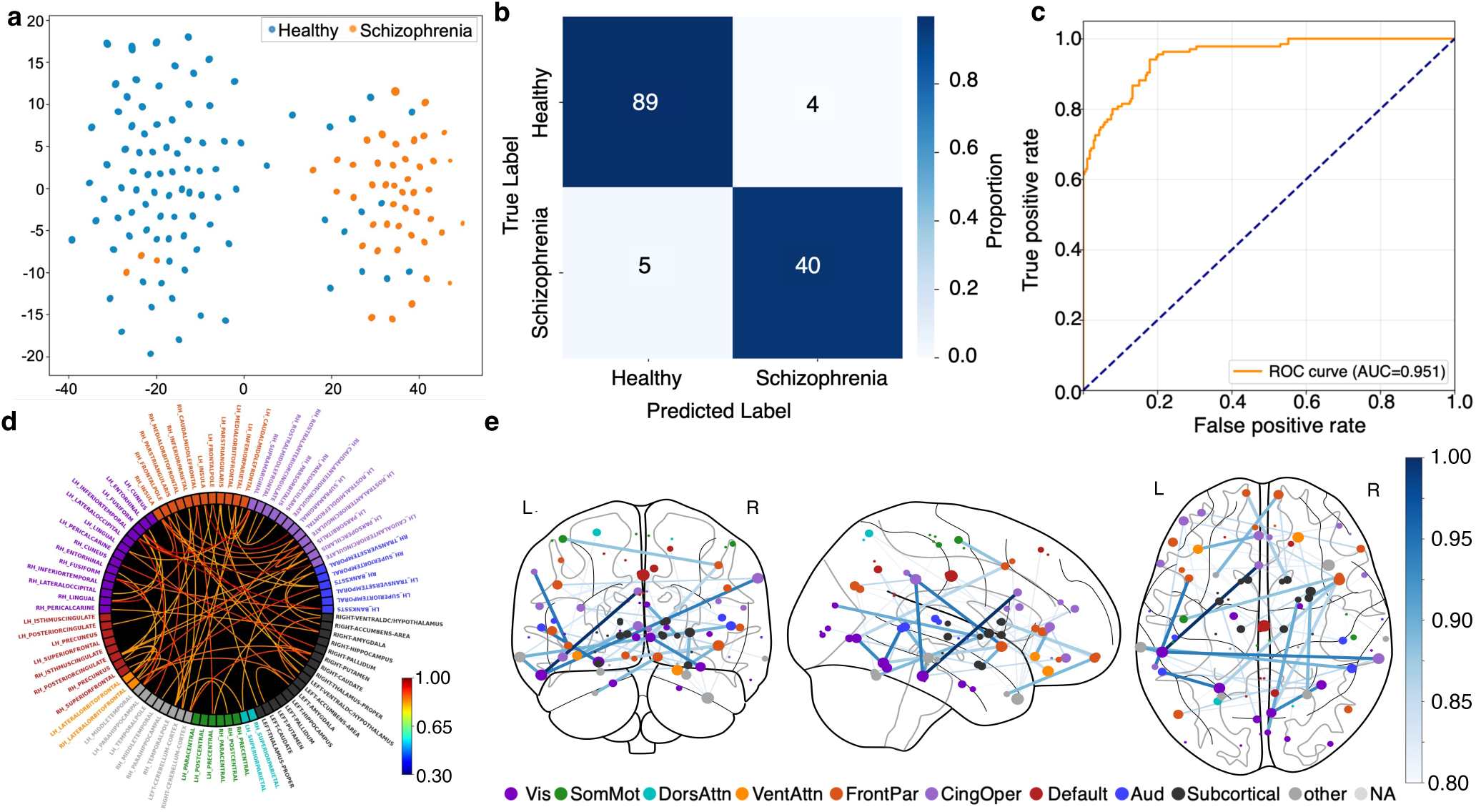
Schizophrenia classification performance and learned connectivity patterns. **a)** t-SNE plot of latent representations for healthy controls and schizophrenia patients. **b)** Confusion matrix of schizophrenia classification. **c)** ROC curve for schizophrenia classification. **d)** Connectivity circle showing functional connections ranked by edge importance (threshold = 0.8), with parcel labels colored by their functional network membership. **e)** Connectome plot projected onto atlas coordinates, showing the spatial distribution of functional connections (threshold = 0.8). Edge thickness is proportional to the learned connection importance score, and node size reflects each region’s node strength computed over the thresholded importance graph.

Edge weights derived from the model are shown in Fig. 7d,e. The model’s classification was based on a specific set of connections including those between the anterior cingulate cortex and inferior temporal gyrus, the supramarginal gyrus and insula, the superior temporal sulcus and fusiform gyrus, and across hemispheres between the right supramarginal gyrus and left inferior temporal gyrus. The connectivity circle (Fig. 7d) highlights the importance of these connections within and between the frontoparietal, cingulo-opercular, auditory, and visual networks, with parcel labels colored by their functional network membership. The connectome map (Fig. 7e) projects these same importance scores onto atlas coordinates, confirming the spatial distribution of these key features. This demonstrates that the model’s high accuracy is driven by identifying disruptions in circuits that are highly consistent with the known pathology of schizophrenia.

#### 2.4.5 Intrinsic functional network analysis of predictive features

To characterize the model’s predictive mechanisms at the level of canonical resting state networks (RSNs), we aggregated the parcel-level importance scores (visualized in Fig. 4b,c, Fig. 5e,f, and Fig. 6c,d) into eight RSNs by normalizing the summed importance weights by network size. This analysis reveals a clear dissociation of predictive features, demonstrating that the functional connections most informative for a given model are highly specific to the predictive target.

As summarized in Figure 8 a, we found that distinct sets of edges are required for three separate tasks. For individual “fingerprinting” (Fig. 8a), the most critical features were high-order connections within and between the control, default mode, and dorsal attention networks, reflecting the unique, intrinsic organization of these systems. In contrast, “task decoding” (Fig. 8b) relied on a different set of edges involved in sensory and motor processing, primarily implicating the visual, somatomotor, and dorsal Attention networks. Finally, “reaction time prediction” (Fig. 8c) was uniquely dependent on a third set of connections, those within or involving subcortical areas related to motor execution and performance monitoring. This clear separation is further quantified by the total network importance (nodal degree of heatmaps in Fig. 8 a-c) shown in Fig. 8e,f. The control, dorsal attention, and default mode network are most critical for fingerprinting, sensory-motor networks dominate for task decoding, and subcortical regions are paramount for predicting reaction time. These results demonstrate that a connection’s “importance” is not a fixed property but is contingent on whether the predictive goal is to identify an individual, a cognitive state, or a behavioral outcome. Furthermore, we examined the features from the multi-task model trained on all three tasks simultaneously (Fig. 8d). This multi-task model learns a composite set of features that forms a superset of the single-task features, leveraging a combination of all the previously identified networks, including default mode, sensory-motor, and subcortical connections to optimize performance across all tasks but has been mostly dominated by the fingerprinting task.

**Figure 8:**
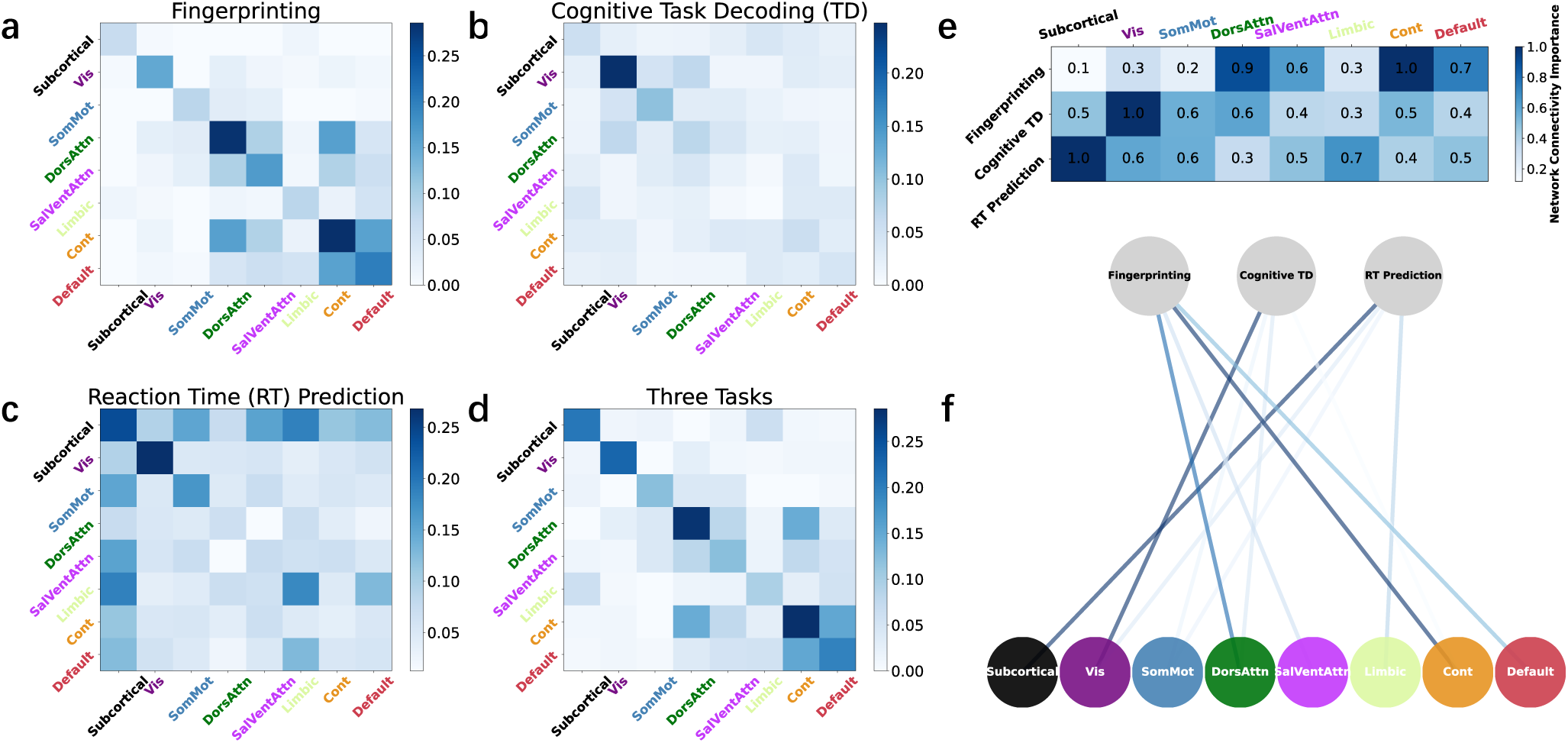
Network-level analysis of predictive connectivity features. **a-d)** Heatmaps of average network-level connectivity importance. These scores were aggregated from parcel-level features (Fig. 4b,c, Fig. 5e,f, and Fig. 6c,d) by calculating the average importance of all constituent parcel-to-parcel edges between any two networks. The heatmaps show the importance for: (a) individual fingerprinting, (b) cognitive task decoding, (c) reaction time prediction, and (d) the multi-task model. **e)** Heatmap of total network importance, quantified as the nodal degree (sum of connections for each network from heatmaps (a-c) and normalized within each task. **f)** Graph representation of the nodal degree data from panel (e).

## 3 Discussion

In this paper, we introduced MAMBAxBrain, a multi-task neural framework that integrates Mamba-based temporal modeling with spatial functional connectivity analysis to address four representative fMRI tasks within a unified architecture. Our results demonstrate that the model achieves state-of-the-art performance in fingerprinting, cognitive decoding, behavioral prediction, and schizophrenia classification, while exhibiting robust cross-session generalization. Importantly, interpretability analyses reveal that MAMBAxBrain does not rely on a diffuse, task-agnostic feature set; instead, it engages biologically distinct circuits for each objective, suggesting that the model captures meaningful structure in the brain’s functional organization rather than superficial statistical regularities. These results shed light on a longstanding debate: rather than relying on either wholly distinct or entirely overlapping systems, the brain appears to preferentially recruit different circuits for identity, cognition, behavior, and pathology while maintaining shared representational structure. Below, we interpret these findings in the context of the brain’s functional organization, compare MAMBAxBrain’s design principles against existing approaches, and consider its potential as a step toward a foundation model for fMRI analysis.

### 3.1 Summary of performance metrics

A fundamental challenge in neuroimaging is the lack of a universal model capable of mining the rich, multidimensional information embedded in fMRI data. While BOLD signals simultaneously encode individual identity, cognitive states, and pathological traits, current analytical frameworks are typically designed to target only one of these dimensions, leaving the others underexploited. In this work, we introduced MAMBAxBrain, a multi-task neural framework designed to bridge this gap by integrating efficient Mamba-based temporal modeling with spatial functional connectivity analysis. This fusion enables a single architecture to achieve superior performance across four representative tasks: fingerprinting, cognitive decoding, behavioral prediction, and psychiatric diagnosis, as shown in Fig. 2 and Table 1. Notably, MAMBAxBrain also exhibited the strongest cross-session generalization, with accuracy reductions of only 1.13% for fingerprinting and 0.54% for cognitive decoding, compared with substantially larger declines of 7.15% and 2.70% for the FST-Mamba model. These results highlight the robustness and generalization ability of MAMBAxBrain across scanning sessions. By jointly modeling temporal dynamics and spatial coordination, MAMBAxBrain demonstrates that these seemingly distinct prediction tasks can be solved within a shared feature space (Fig. 3). The model achieved an *R*^2^ of 0.72 in reaction time prediction without training on a large dataset, suggesting that fingerprinting and cognitive decoding tasks share transferable behavior-related representations. Because fingerprinting and cognitive decoding are trained on the same input data and differ only in their labels, this improvement cannot be explained by additional training data; it reflects genuine cross-task transfer from the task-decoding objective.

### 3.2 Neuroscientific insights revealed by MAMBAxBrain

Beyond achieving high accuracy, our interpretability analyses reveal that MAMBAxBrain solves diverse neuroimaging tasks by exploiting the brain’s intrinsic functional hierarchy. Rather than relying on a diffuse, “one-size-fits-all” feature set, the model disentangles distinct circuit mechanisms that align with the specific biological demands of each problem. We emphasize that these findings are model-derived attributions obtained with Integrated Gradients rather than direct measures of neural causation; the network-level patterns highlighted below should therefore be read as the connections most informative to the model for each task, to be corroborated by future targeted experimental and lesion-based studies.

#### 3.2.1 Functional fingerprinting

For individual identification, the model preferentially attended to higher-order association cortices, specifically the control, default mode, and dorsal attention networks (Fig. 8a). This aligns with the functional fingerprint hypothesis (Finn et al., 2015), which suggests that evolutionarily recent, highly variable transmodal hubs carry the unique signature of an individual’s identity. Further, the fact that model performance on cross-session generalization was the highest for resting fMRI suggests that the most stable and informative representations of identity are present at rest in comparison to task states.

#### 3.2.2 Cognitive task decoding

In striking contrast, cognitive state decoding relied on primary sensory, motor, and ventral attention networks (Fig. 8b). This double dissociation mirrors recent large-scale findings (Hannum et al., 2023) and highlights a fundamental trade-off: while the distinctive architecture of higher-order networks defines *who* a person is, the stereotyped engagement of sensory-motor systems defines *what* they are doing.

#### 3.2.3 Behavioral prediction

Moving beyond cortical topography, the model demonstrated that behavioral speed (cognitive task reaction time) is governed by a deeper, subcortical circuitry (Fig. 8c). Prediction of millisecond-scale variability was driven by subcortical-motor and subcortical-limbic loops. Distinct from the cortical attention networks emphasized in previous predictive modeling studies (Yoo et al., 2018), this circuitry maps onto the classical motor execution pathway (subcortical to somatomotor) where the basal ganglia and thalamus initiate motor commands (Mink, 1996). Crucially, this circuitry also included subcortical to ventral attention connectivity, reflecting the detection of salient cues (Corbetta & Shulman, 2002), and leveraged the limbic system, suggesting that processing speed is modulated by internal arousal and motivational states (Pessoa, 2009).

#### 3.2.4 Schizophrenia classification

Finally, in the context of pathology (schizophrenia), MAMBAxBrain identified a specific breakdown in the interaction between these hierarchical levels. Diagnosis was driven by dysconnectivity in top-down control circuits, specifically linking the anterior cingulate cortex and insula to auditory and visual association regions (Fig. 7d,e). This finding is consistent with the dysconnection hypothesis (Friston, 1998; Stephan et al., 2009), pointing to a failure of control networks to constrain sensory processing streams. The disconnection of salience hubs (anterior cingulate cortex, insula) from the superior temporal sulcus and fusiform gyrus aligns with theories of aberrant salience and a “blurring” of internal and external reality (Menon, 2011), while specific deficits in superior temporal sulcus and supramarginal gyrus connectivity map onto the hallmark social and linguistic symptoms of the disorder (DeLisi, 2001; Green et al., 2015; Palaniyappan & Liddle, 2012; Palaniyappan et al., 2012).

### 3.3 Analysis of baseline models

To better understand the performance differences among baseline models, we summarize and discuss the competing approaches from the perspectives of input feature and model architecture. SPDNet, BrainNetTF, and CorrNN follow an FC-only paradigm: they take functional connectivity matrices (or correlation-derived features) as input and focus on modeling static or aggregated connectivity patterns. Specifically, SPDNet emphasizes symmetric positive definite (SPD) representations and Riemannian geometric modeling; BrainNetTF adopts a Transformer architecture to capture relational structure in brain networks; and CorrNN performs discriminative learning based on connectivity features. A potential limitation of these methods is that when a task strongly depends on temporal dynamics (e.g., subject-specific variability across sessions or behavior-related dynamics), relying solely on FC may attenuate information about temporal order and moment-to-moment fluctuations, thereby limiting generalization. Moreover, performance can be sensitive to the FC construction strategy and measurement noise. Mamba follows a TS-only paradigm: it efficiently models long-range temporal dependencies via a state-space model and is appropriate for extracting dynamic patterns directly from raw multiregional time series. However, cross-region interactions are largely learned implicitly without explicit network-structure constraints, which may result in less robust representations under session shifts or in settings where stable connectivity priors are beneficial. FST-Mamba and MAMBAxBrain both fall within the combined FC–TS paradigm, aiming to leverage temporal dynamics alongside network-level information. FST-Mamba models temporal information indirectly through sequences of dynamic functional network connectivity (dFNC) rather than directly from raw BOLD signals, emphasizing evolving inter-regional interactions but potentially obscuring fine-grained signal-level temporal structure. Moreover, its complex hierarchical architecture incurs high computational cost and complicates optimization, which negatively impacts performance. FST-Mamba is evaluated using separate single-task training, whereas MAMBAxBrain is designed to learn shared representations that support multi-task learning. Implementation details for all baseline models are provided in Supplementary Material.

### 3.4 Implications and potential applications of MAMBAxBrain

Collectively, these results demonstrate that MAMBAxBrain does not merely learn statistical correlations, but successfully maps the diverse dimensions of brain function, identity, cognition, behavior, and pathology onto their biologically relevant set within a multi-task feature space. Many neuroscientific insights may emerge from these findings. Historically, it has been challenging to uncover the neural correlates of behavior, since behavioral signatures shared across individuals are frequently masked by individual-specific neural patterns. That individual identity is subserved by higher-order association cortices (DMN, Frontoparietal), whereas task-specific mental activity is driven by sensory-motor systems, sheds new light on this question and merits further exploration. “Reaction time prediction” was uniquely dependent on a third set of connections, those within or involving subcortical areas related to motor execution and performance monitoring. Intriguingly, identity-related connections were dominantly ipsilateral, while task-related connections were strikingly inter-hemispheric, suggesting that cognitive tasks require wide coordination between sensorimotor areas across hemispheres, while identity-related concepts are more lateralized. More generally, our work effectively delineates representational structures responding to cognitive tasks that are shared across individuals, versus idiosyncratic spatial structures that are tuned to identity. Although these concepts have been broadly surmised in the literature, our use of a deep learning framework introduces a quantitative and computational aspect that was hitherto missing.

Nevertheless, these subsystems all work together for any given task domain. MAMBAxBrain was able to train on all tasks simultaneously and did not incur an appreciable performance penalty vis a vis single-task training. That seemingly distinct tasks undertaken by the brain can be solved within a shared representational space, suggests the existence of transferable, behavior-related representations in the human brain. The flexibility of MAMBAxBrain enables a wide range of applications across cognitive neuroscience, behavioral prediction, and clinical research. Its dual-stream backbone learns task-agnostic representations, while the multi-task heads enable those representations to be adapted efficiently to new tasks. For cognitive neuroscience, the model could support fine-grained decoding of mental states, mapping of task-evoked representations, and examination of individual differences in cognition. For behavioral prediction, the framework may enable accurate forecasting of reaction time, vigilance, learning trajectories, and other dynamic behaviors that unfold over time. In clinical contexts, MAMBAxBrain could facilitate early detection of neuropsychiatric and neurodegenerative conditions, patient stratification based on functional signatures, and monitoring of treatment response. Beyond these dimensions, the model may also serve as a general tool for harmonizing heterogeneous datasets, transferring knowledge across cohorts, and building robust biomarkers that generalize across scanners, populations, and acquisition protocols.

### 3.5 Future directions toward more generalizable fMRI models

MAMBAxBrain exhibits several properties that are aligned with goals of more generalizable or foundation models for fMRI analysis. In particular, it shows robustness across sessions, scalability to long temporal sequences, and the ability to support multiple tasks within a unified architecture. These characteristics suggest its potential to move beyond task-specific pipelines toward more flexible and reusable representations. Future work will explore architectural extensions that enable scalable specialization, such as mixture-of-experts mechanisms, as well as large-scale self-supervised pretraining and improved cross-site harmonization. The development of transferable, task-agnostic representations remains an open challenge, and MAMBAxBrain can be viewed as a step in this direction.

### 3.6 Limitations

While MAMBAxBrain demonstrates robust performance and strong cross-task generalization in multiple cognitive and clinical dimensions, several aspects warrant further investigation. First, the schizophrenia dataset used in this study has a relatively small sample size, which may limit the generalizability of results to broader clinical populations. Consequently, under diagnosis-stratified, subject-disjoint 5-fold cross-validation the performance is both lower and considerably more variable than a single within-split estimate (accuracy 82.14%±11.03%, AUC 0.83±0.17; best among all models, Supplementary Table S6), and on the independent COBRE cohort the model generalized above chance but with reduced sensitivity (AUC 0.81, accuracy 0.68; Supplementary Table S7). These results should therefore be regarded as preliminary, and larger, multi-site clinical samples will be needed to establish robust diagnostic performance. Second, despite its robust cross-session generalization and the encouraging external schizophrenia result on COBRE, systematic evaluation across additional independent cohorts and imaging sites remains an important next step, particularly for the HCP-based fingerprinting, decoding, and reaction-time tasks, which were not externally validated here. Third, although MAMBAxBrain attains a high overall *R*^2^ of 0.92 for reaction-time prediction, this largely reflects systematic differences in mean RT between conditions. As reported in Supplementary Table S3, the model explains substantially less variance within individual conditions (mean within-condition *R*^2^ ≈ 0.19), indicating that individual within-condition RT prediction remains limited and is an important target for future work. Finally, the framework currently relies on atlas-based cortical parcellation, which facilitates standardized feature extraction and interpretability within individual datasets. However, this constrains multi-task training across datasets with heterogeneous parcellations, as regional definitions and boundaries are not directly comparable. This parcellation difference also bears on the cross-dataset interpretability comparison. Because the Desikan–Killiany atlas follows gyral anatomy whereas the Schaefer atlas is derived from functional gradients, individual ROI–ROI importance values are not directly comparable between the UCSF schizophrenia analysis and the HCP-based analyses. For this reason we report interpretability at the level of canonical resting-state networks rather than individual parcels: both atlases map onto the same small set of networks (e.g., frontoparietal/control, cingulo-opercular, default mode, and sensorimotor systems), and aggregating edge importance to the network level mitigates the parcellation mismatch and supports the qualitative cross-task and cross-dataset comparisons reported here. A fully quantitative ROI-level comparison across atlases would require resampling to a common parcellation space, which we leave to future work alongside atlas-agnostic harmonization strategies for joint multi-task training. Note that the interpretability maps should not be viewed as absolute biological ground truth or direct causal pathways. Instead, they represent model-derived feature attributions indicating which macroscopic functional connectivity patterns the architecture prioritizes during prediction.

## 4 Methods

### 4.1 fMRI data acquisition and preprocessing

This study utilized data from multiple sources. The primary normative cohort was drawn from the Human Connectome Project (HCP) Young Adult S1200 release (Van Essen et al., 2013), Test-Retest (Van Essen et al., 2012), and a separate schizophrenia cohort collected at UCSF. The detailed dataset descriptions, including subject counts, ROI numbers, and data sources, are provided in the Supplementary Table S1.

#### 4.1.1 HCP rest and task dataset

We used 770 subjects from the Human Connectome Project Young Adult (HCP-YA) dataset (Van Essen et al., 2013) in this study. We excluded all HCP subjects that had documented quality control issues. Unprocessed HCP data were downloaded and organized into BIDS format. Anatomical scans were collected with an MPRAGE T1-weighted sequence. Functional MRI data included four 15-minute rest sessions and seven task sessions (Working Memory, Language, Motor, Social, Relational, Gambling, Emotion), all with a repetition time (TR) of 720 ms. We applied a state-of-the-art image processing pipeline, micapipe (Cruces et al., 2022), which processes the anatomical, functional, and diffusion MRI data in a coherent framework to produce subject-specific structural connectivity and functional regional time series in the Schaefer atlas with 100 cortical regions (Schaefer et al., 2018), combined with 14 subcortical regions from the FreeSurfer segmentation (Patenaude et al., 2011). The fMRI underwent image re-orientation, motion and distortion correction, and nuisance signal regression (white matter, cerebrospinal fluid (CSF), and frame-wise displacement spikes). Volumetric timeseries were mapped to the native Freesurfer space using boundary-based registration (Greve & Fischl, 2009). Vertex-level timeseries were averaged into parcels defined by the Schaefer atlas with either 100 cortical regions (Schaefer et al., 2018). Parcel-level timeseries were bandpass filtered to be less than 0.1Hz.

To assess the reliability of our derived measures, we also utilized the Human Connectome Project Test-Retest (HCP-RT) dataset (Van Essen et al., 2012). This dataset consists of 45 participants from the main cohort who returned for a second full scanning session, identical to the first, approximately 4-6 months later. This longitudinal data allows for the evaluation of within-subject stability of the functional and structural connectivity metrics. To investigate the neural predictors of behavioral outcomes across different cognitive tasks, we extracted specific reaction time (RT) measures from the HCP task battery. For each subject, we used the mean RT (averaged across the whole runs) from the Working Memory, Relational, and Language (both Math and Story components) tasks. From the Social task, we utilized RT from both the ‘Theory of Mind’ and ‘Random’ conditions (Barch et al., 2013).

#### 4.1.2 Schizophrenia dataset

The clinical data were collected as part of a multisite study in collaboration with the Brain Imaging and EEG Laboratory at the San Francisco VA Medical Center. This cohort consisted of 45 patients with schizophrenia and 93 healthy controls. Scanning was conducted at the UCSF Neuroimaging Center on a Siemens 3T TIM TRIO scanner. High-resolution T1-weighted structural images were acquired using an MPRAGE sequence (TR = 2300 ms; TE = 2.95 ms; flip angle = 9°; FOV = 256×256; slice thickness = 1.20 mm). Resting-state fMRI data were collected using a T2*-weighted echo-planar imaging (EPI) sequence (TR = 2000 ms; TE = 29 ms; flip angle = 75°; 191 volumes).

The T1-weighted and BOLD images for the clinical cohort were preprocessed using fMRIPrep (Esteban et al., 2019), a robust pipeline based on Nipype (Gorgolewski et al., 2011) and NiLearn (Abraham et al., 2014). For more details on this pipeline, see the section corresponding to workflows in fMRIPrep’s documentation. For completeness, we summarize the anatomical and functional preprocessing below. T1-weighted (T1w) images were corrected for nonuniformity (Avants et al., 2008; Tustison et al., 2010) and were used as a reference image throughout the workflow. The T1w reference was skull stripped and segmented based on cerebrospinal fluid, white matter, and gray matter (Avants et al., 2008; Zhang et al., 2002). The T1w reference and template were spatially normalized to a standard space (MNI152NLin2009cAsym) with nonlinear registration (Avants et al., 2008). FMRIPrep generated a BOLD reference image that was coregistered to the T1w reference using flirt and boundary-based registration (Greve & Fischl, 2009; Jenkinson & Smith, 2001). Head-motion parameters were estimated before spatiotemporal filtering using FSL’s mcflirt (Jenkinson et al., 2002). BOLD runs were slicetime corrected and resampled onto their original, native space by applying transforms to correct for head motion (Cox & Hyde, 1997). The BOLD time series were resampled into the same standard space as the T1w image (MNI152NLin2009cAsym).

The whole brain was parcellated into 86 regions of interest (68 cortical, 18 subcortical) as defined by the Desikan–Killiany atlas (Desikan et al., 2006), from which average functional time series were extracted. To denoise the functional data, we implemented the ICA-AROMA with Global Signal Regression (GSR) pipeline, following the procedures benchmarked in (Ciric et al., 2017). The first stage of this pipeline, ICA-AROMA, performed automatic detection and removal of signal noise components, including those driven by head motion (Pruim et al., 2015). Subsequently, GSR was applied to remove the mean, brain-wide signal at each time point. The pipeline included smoothing with a 6 mm FWHM SUSAN kernel and regional time series bandpass filtering with a Butterworth filter from 0.01 to 0.25 Hz.

To directly address the possibility of within-subject information leakage and to characterize the variability of the schizophrenia result, the UCSF cohort was additionally evaluated with diagnosis-stratified, subject-disjoint 5-fold cross-validation, in which all windows of a given subject were assigned entirely to a single fold so that no subject appeared in both the training and test sets, and the patient/control ratio was preserved across folds. To further assess generalization beyond the UCSF cohort, we evaluated the trained model on the independent, publicly available COBRE schizophrenia dataset (72 patients and 74 healthy controls) as an external test set, with no COBRE data used for training; external performance is reported in Supplementary Table S7.

The COBRE dataset (Aine et al., 2017) was obtained from the NIAK-preprocessed release (Bellec, 2015) available on figshare, itself a derivative of the original COBRE sample distributed through the International Neuroimaging Data-sharing Initiative (INDI). This release comprises resting-state fMRI data from 72 patients with schizophrenia (58 male; age range 18–65 years) and 74 healthy controls (51 male; age range 18–65 years), each with up to 150 EPI BOLD volumes (TR = 2000 ms, TE = 29 ms, flip angle = 75°, 32 slices, voxel size 3×3×4 mm^3^, matrix size 64×64) acquired in MNI152 non-linear symmetric space at 3 mm isotropic resolution. We used the full released cohort (72 patients, 74 controls). Preprocessing was performed using the NeuroImaging Analysis Kit (NIAK v0.12.14) and included slice-timing correction, rigid-body motion correction, coregistration to each subject’s T1-weighted scan, and non-linear registration to MNI space via the CIVET pipeline; nuisance regression removed slow time drifts, white-matter and ventricular signals, and motion-derived confounds, and volumes were spatially smoothed with a 6 mm isotropic Gaussian kernel. To match the feature space used for the UCSF cohort, regional time series were subsequently extracted by parcellating the preprocessed volumes into the same 86-region Desikan–Killiany atlas applied to the UCSF data.

#### 4.1.3 Sliding-window sample generation

To ensure consistent modeling across heterogeneous datasets, MAMBAxBrain adopts a multi-task strategy that converts fMRI time series into fixed-shape, multichannel temporal tensors. This allows all data to be processed by a common model backbone for representation learning. Given a time series of length *T*, sliding window of length *L* and stride *δ* are applied along the temporal axis to generate *N* samples as follows:

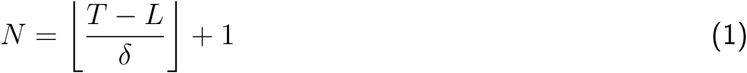

Each sample contains *L* consecutive time points, representing a localized brain state. By adjusting the window length and stride, this procedure flexibly controls temporal resolution and overlap, enabling adaptation to diverse experimental designs and modeling requirements. After sample generation, if sequence lengths vary across subjects or sessions, a zero-padding operation is applied to align all samples to a uniform length, ensuring consistent tensor shapes during batch processing. Subsequently, all time series are z-score normalized on a per-ROI basis, where each ROI’s time series is standardized using its own mean and standard deviation across the entire sequence to remove baseline differences among subjects and brain regions. In this way, different tasks are seamlessly integrated by simply adjusting the sliding-window sampling parameters to the desired configuration.

### 4.2 MAMBAxBrain

#### 4.2.1 Framework overview

MAMBAxBrain is a multi-task architecture for large-scale modeling of neural dynamics from multiregional fMRI time series. Existing deep fMRI models typically specialize in either spatial modeling of connectivity patterns between regions or focusing on time-series dynamics within regions, but rarely integrate both within a spatiotemporal computational framework. As illustrated in Fig. 1, MAMBAxBrain addresses this limitation by coupling two components: a temporal signal stream *T*(·) based on the established Mamba Selective State Space Model (SSM) (Gu & Dao, 2024), which captures long-range temporal dependencies; and a newly introduced functional connectivity (FC) stream *S*(·) that explicitly models inter-regional relationships from pairwise connectivity matrices. While prior Mamba-based architectures have primarily focused on temporal modeling of neural signals, MAMBAxBrain extends this framework by integrating FC-based spatial modeling and introducing a learnable fusion mechanism that jointly encodes temporal and connectivity information. This dual-stream and fusion design constitutes the core methodological innovation of MAMBAxBrain, enabling unified modeling of temporal and connectivity representations within a single, generalizable architecture. Their latent embeddings are fused through a feature integration module *F*(·), yielding a unified representation **z** that serves as a common feature space for downstream tasks.

Unlike existing task-specific networks (Gu & Dao, 2024; Huang & Van Gool, 2017; Kan et al., 2022; Wei et al., 2025), MAMBAxBrain maintains a single, generalizable backbone. By attaching lightweight task-specific heads to the shared representation **z**, the framework enables scalable fMRI representation learning across diverse tasks without redesigning the network. This modular structure supports a broad range of objectives, including brain fingerprinting, cognitive decoding, behavioral prediction, and clinical diagnosis. Formally, for an input sample **X** ∈ ℝ*^L^*^×^*^D^*, where *L* is the number of time points and *D* is the number of brain regions, the framework is defined as:

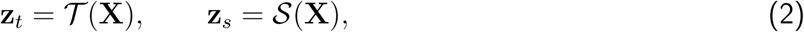

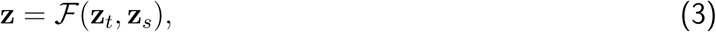

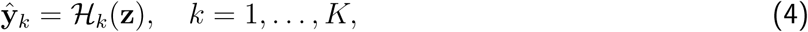

where **z***_t_* and **z***_s_* denote the embeddings from the temporal and FC streams, respectively, and **z** is the fused, task-agnostic representation. Each task head *H_k_*(·) generates the prediction **ŷ***_k_* for the *k*-th task.

#### 4.2.2 Temporal signal stream

In neural signal modeling, the temporal dimension captures the activity pattern of each brain region over time, which is essential for building effective neural representations. MAMBAxBrain uses the Selective State Space Model (Gu & Dao, 2024) to capture long-range temporal dependencies in fMRI sequences. The selective SSM dynamically modulates temporal information, attenuates noise, and highlights salient time points, thereby enhancing robustness when modeling fMRI data with inherently low signal-to-noise ratios. In addition, the linear state update mechanism ensures the computational complexity scales linearly with sequence length, which is substantially more efficient than self-attention for long-sequence modeling.

Each sampled fMRI segment (**X** ∈ ℝ*^L^*^×^*^D^*) first passes through a one-dimensional convolution layer to capture local temporal signal variations, and is subsequently processed by the selective SSM to extract long-range temporal dependencies. The evolution of neural states is represented by a hidden state vector **h***_t_*, which is recursively updated along the temporal dimension as:

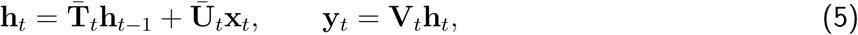

where **T̅***_t_* ∈ ℝ*^N^*^×^*^N^* is the discretized state transition matrix that controls how hidden states propagate and retain memory across time steps; **U̅***_t_* ∈ ℝ*^N^*^×^*^D^* is the discretized input projection matrix that determines how the current brain-region input affects state updates; **V***_t_* ∈ ℝ*^D^*^×^*^N^* is the output projection matrix that maps hidden states back to the observable space; *N* denotes the dimensionality of the hidden state vector. To improve adaptability across varying temporal contexts, these matrices are parameterized as input-dependent functions, allowing the model to dynamically adjust its update rate and information-retention strength for selective temporal propagation and filtering.

To maintain stable gradient propagation during temporal updates, residual connections are incorporated in the module to preserve information flow and alleviate the vanishing gradient problem, while normalization layers further stabilize the training dynamics. Next, all hidden states are aggregated along the temporal dimension, and temporal mean pooling is used to compress them into a compact embedding vector:

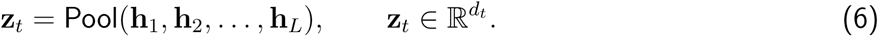

Here, Pool(·) denotes the global mean pooling operation along time, **z***_t_* is the final temporal embedding vector, and *d_t_* is its dimension. This resulting embedding of the temporal signal stream is projected into a shared representation space and fused with the FC stream to achieve joint spatiotemporal modeling.

#### 4.2.3 Functional connectivity stream

In neural signal modeling, inter-regional relationships are a key determinant of brain function. Therefore, MAMBAxBrain incorporates a functional connectivity stream dedicated to learning spatial relationships among brain regions. The input to the FC stream is identical to that of the temporal stream, a sampled fMRI time-series segment **X** ∈ ℝ*^L^*^×^*^D^*. To quantify the functional association between brain regions, a mapping function, denoted as Φ(·), is applied to the time-series segment **X**. This yields a symmetric functional connectivity matrix **C**:

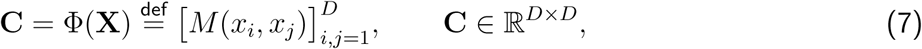

where *x_i_, x_j_* ∈ ℝ*^L^* are the BOLD time series of brain regions *i* and *j*, and *M* (·) is a pairwise association measure, such as Pearson correlation or covariance.

The resulting symmetric matrix **C** is vectorized by extracting its upper triangular elements (excluding the diagonal) and flattening them into a vector **v** ∈ ℝ*^E^*, where 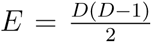 is the number of unique inter-regional connections. A connectivity encoder, consisting of multilayer perceptrons (MLP), maps the high-dimensional connectivity vector into a compact, task-agnostic spatial representation:

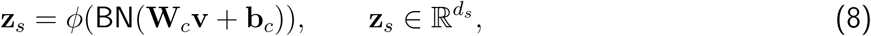

where **W***_c_* and **b***_c_* are learnable parameters, BN denotes batch normalization, and *ϕ*(·) is a nonlinear activation function. The resulting spatial embedding vector **z***_s_* is then projected into the shared feature space.

#### 4.2.4 Feature fusion

While temporal and FC embeddings capture complementary aspects of brain dynamics, effective integration of these heterogeneous representations is essential for multi-task modeling. MAMBAxBrain employs a learnable feature fusion module that adaptively integrates temporal and spatial information within a shared latent space. The temporal embedding 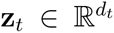 and the FC embedding 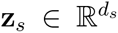 are first concatenated along the feature dimension:

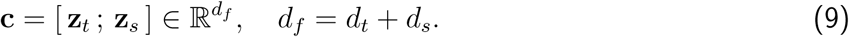

To enable nonlinear interaction and cross-modal alignment between temporal and FC features, the concatenated feature **c** is processed by a fully connected layer, nonlinear activation, and dropout regulariza-tion:

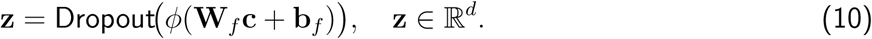

Here, 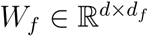 and **b***_f_* are learnable parameters. This fusion mechanism yields a more expressive and adaptive spatiotemporal representation. The resulting fused feature **z** serves as a shared representation for downstream tasks.

#### 4.2.5 Task heads

After obtaining the fused representation **z** ∈ ℝ*^d^*, a set of lightweight task heads is designed to project it into the output space of each task:

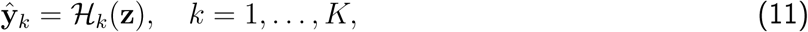

where *H_k_*(·) denotes the prediction head for the *k*-th task. The current implementation includes four task heads corresponding to brain fingerprinting, cognitive task decoding, behavioral reaction time prediction, and schizophrenia classification. Each task head is designed as a lightweight module, typically a linear layer with negligible parameter count relative to the backbone network. For classification tasks (brain fingerprinting, cognitive task decoding, and schizophrenia classification), the head outputs a probability vector over the corresponding classes. For the regression task (reaction time prediction), it produces a continuous value reflecting individual behavioral or response characteristics.

This modular architecture decouples task-specific objectives from the shared embedding, allowing the backbone network to remain fixed while new tasks can be incorporated seamlessly by attaching additional heads. Such a design promotes scalability, multi-task generalization, and efficient expansion across cognitive and clinical dimensions.

### 4.3 Experimental settings

All experiments were implemented in PyTorch and executed on a single NVIDIA RTX 3080 Ti GPU. To ensure robustness, each experiment was repeated three times with different random seeds, and early stopping was applied to prevent overfitting. A comparison of computational cost (parameters, FLOPs, and training/inference time) against all baselines, measured under matched inputs and a common state dimension on this hardware, is reported in Supplementary Table S5. Training configurations are summarized in Table 2. We started from the default hyperparameters of the Mamba model (Gu & Dao, 2024) and performed limited fine-tuning using cross-validation to ensure stable training. The final settings were adjusted according to the characteristics of each task to achieve a balance between model capacity and overfitting risk. In particular, tasks with longer temporal windows adopted larger state dimensions to better capture long-range dependencies, while maintaining computational efficiency across all experiments.

**Table 2:** Training configurations across all tasks. “Varied” indicates that the values vary across tasks during joint multi-task training. All entries are MAMBAxBrain’s own per-task configurations rather than a comparison across models; the larger state dimensions accompany the tasks with longer temporal windows (e.g., reaction time prediction, window 464) to capture longer-range dependencies, whereas the short-window classification tasks use smaller values. The split column reports the primary protocol used for cross-model comparison in Table 1; the schizophrenia task is additionally evaluated with diagnosis-stratified, subject-disjoint 5-fold cross-validation reported in the text, and the HCP within-subject tasks use temporal 5-fold cross-validation as a robustness check.

| Configuration | Brain Fingerprinting | Cognitive Task Decoding | Reaction Time Prediction | Schizophrenia Classification | Multi-Task Training |
| --- | --- | --- | --- | --- | --- |
| Window Size | 100 | 100 | 464 | 70 | Varied |
| Total Samples | 3,431k | 3,431k | 4,347 | 5,796 | 3,435k |
| Train/Test Split | 7:3 | 7:3 | 7:3 | 6:4 | 7:3 |
| State Dimension | 16 | 16 | 150 | 2 | 150 |
| Kernel Size | 4 | 4 | 16 | 2 | 16 |
| Expand Ratio | 3 | 3 | 16 | 1 | 16 |
| Optimizer | AdamW | AdamW | AdamW | AdamW | AdamW |
| Learning Rate | $8 \times 10^{-4}$ | $8 \times 10^{-4}$ | $1 \times 10^{-4}$ | $8 \times 10^{-4}$ | $5 \times 10^{-4}$ |
| Batch Size | 128 | 128 | 128 | 128 | 128 |
| Epochs | 60 | 60 | 60 | 60 | 60 |

In single-task experiments, brain fingerprinting, cognitive task decoding, and reaction time prediction were performed using the HCP–Young Adult dataset (770 subjects, 18 scans per subject, 114 ROIs).

The schizophrenia classification task used a clinical dataset collected at UCSF, comprising 93 healthy controls and 45 patients with 86 ROIs. To prevent data leakage, all classification tasks were split into training and validation sets before applying the sliding-window procedure. The split was performed on the raw ROI time series before any window was extracted, so that no window spans the train/test boundary and training and test windows never share a time point. The split unit was task-dependent: for brain fingerprinting and cognitive task decoding it was a within-subject temporal segment of each scan, since both tasks require the same subjects to be present in training and test (fingerprinting is a 770-way individual-identification task), whereas reaction time prediction was split at the level of individual (subject × condition) regression samples. Cross-subject generalization was assessed separately on the independent cross-session HCP-RT data. The main tables report the primary split listed in Table 2 (7:3 for the HCP within-subject tasks) as mean ± standard deviation across three random seeds. As an additional robustness check for the within-subject tasks, we also used temporal 5-fold cross-validation: each scan’s time series was divided into five contiguous, non-overlapping blocks that tile the entire recording, and each block served once as the held-out test segment while the remaining blocks formed the training segment, with performance averaged across the five folds. The window stride was fixed at 1, and window lengths followed the task-specific configurations listed in Table 2. Because a stride of 1 yields overlapping windows, we additionally evaluated fingerprinting and cognitive decoding with non-overlapping windows (stride equal to the window length) as a sensitivity analysis; results are reported in Supplementary Table S2.

For reaction time prediction, sequences were zero-padded to achieve uniform temporal length. Mean absolute error (MAE) was used as the loss function for regression, and cross-entropy for classification. The cross-session experimental settings were identical to those used in the single-task experiments. The details of hyperparameter tuning for our model can be found in the Supplementary Material (Implementation details and hyperparameter setting).

In the multi-task training experiments, three configurations were designed to systematically evaluate the model’s scalability when incorporating new tasks. The brain fingerprinting task served as the base task for learning generalized individual-level functional connectivity representations. First, the model was trained to obtain stable subject embeddings and then extended to cognitive task decoding and reaction time prediction to assess scalability under limited-data conditions. Next, joint training on brain fingerprinting and cognitive task decoding was conducted to examine its ability to learn shared representations across classification tasks and to evaluate scalability on reaction time prediction. Finally, all three tasks were trained jointly to assess overall representation learning and inter-task complementarity. For all adaptation experiments, 20% of the target task data was used for fine-tuning.

A multi-data loader training strategy was employed to coordinate updates across different tasks and maintain balanced learning dynamics. During training, batches from task-specific datasets were alternated according to the schedule: the model trained on two classification batches (brain fingerprinting and cognitive task decoding), followed by one regression batch (reaction time prediction), to ensure balanced optimization across task types. The overall loss function for multi-task training was formulated as a weighted composite loss:

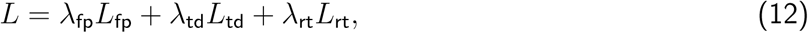

where *L*_fp_, *L*_td_, and *L*_rt_ represent the task-specific losses for brain fingerprinting, cognitive task decoding, and reaction time prediction, respectively, and *λ*_fp_, *λ*_td_, and *λ*_rt_ are their corresponding weights. Because regression tasks are generally more difficult to optimize, a larger weight was assigned to the reaction time prediction loss to ensure balanced learning across tasks.

### 4.4 Feature importance analysis

To interpret the key connectivity patterns leveraged by the model in brain fingerprinting and related tasks, we quantify the contribution of each functional connection to the model’s prediction using Integrated Gradients (Sundararajan et al., 2017), a standard axiomatic attribution method, and map the resulting scores onto the ROI–ROI connection space to obtain an interpretable connectivity importance map. Concretely, for each subject we define an all-zero connectivity baseline and linearly interpolate from this baseline to the observed functional connectivity; the attribution of each connection is the product of its deviation from the baseline and the gradient of the model output with respect to that connection, averaged along the interpolation path, with the time-series input held at its observed value. These attributions satisfy the completeness axiom, summing to the difference in model output between the observed connectivity and the baseline, and we aggregate their absolute values across subjects to obtain a stable importance score for each ROI–ROI connection. The scores are then reshaped into a symmetric ROI×ROI matrix, min–max normalized to [0, 1], and projected onto atlas coordinates for visualization.

## Data and Code Availability

All data used in this study are publicly available from the Human Connectome Project and the UCSF Schizophrenia dataset. All source code for the MAMBAxBrain framework will be made publicly available on GitHub upon publication.

## Author Contributions

Y.X. and F.A. developed and evaluated the machine learning framework and co-wrote the manuscript. B.S. ran the preprocessing pipeline. Y.X. carried out model training, testing, and benchmarking. U.S., B.S., and G.G. contributed to the results discussion and manuscript revision. A.R. and M.C. conceived and supervised the project, and revised the manuscript. All authors discussed the results and commented on the manuscript throughout all stages of the work.

## Acknowledgments

The authors thank the Human Connectome Project and UCSF Neuroimaging Center for data access and technical support.

