## Supplementary Material for "MAMBAxBrain: A Multi-task Neural Framework Linking Brain Functional Dynamics to Individual Fingerprints, Cognitive and Disease States"

### Supplementary Material for "MAMBAxBrain: A Unified Neural Framework Linking Brain Functional Dynamics to Individual Fingerprints, Cognitive and Disease States"

#### Dataset description

Table S1 summarizes the dataset characteristics used across all tasks, including data sources, number of subjects, total scans, and ROI definitions. HCP datasets were used for brain fingerprinting, cognitive task decoding, and reaction time prediction tasks, while the UCSF dataset was used for schizophrenia classification. For multi-task training, all HCP-based tasks were jointly included to enable shared representation learning across domains.

Table S1: **Dataset characteristics across all tasks.**

| **Configuration** | **Brain Fingerprinting** | **Cognitive Task Decoding** | **Reaction Time Prediction** | **Schizophrenia Classification** | **Multi-Task Training** |
|---|---|---|---|---|---|
| Data Source | HCP | HCP | HCP | UCSF | HCP |
| Subject Number | 770 | 770 | 770 | 138 | 770 |
| Total Scans | 5,998 | 5,998 | 4,347 | 138 | 10,345 |
| ROI Count | 114 | 114 | 114 | 86 | 114 |

#### Implementation details and hyperparameter setting

For the parameter selection of MAMBAxBrain, we conducted systematic tuning. For the temporal modeling component, we started from the default configuration of the Mamba model and performed a limited grid search over key hyperparameters, such as the state dimension, kernel size, and expansion ratio, to adapt the model to different task-specific temporal window lengths and signal characteristics. For the functional connectivity modeling component, we also performed hyperparameter search, mainly adjusting the number of layers, hidden dimensions, and activation functions of the multilayer perceptron to balance model capacity and generalization ability. All experiments were conducted under the same hardware environment, and each task was repeated three times to evaluate performance stability.

For all benchmark models (SPDNet, Brain Network Transformer, Mamba, and FST-Mamba), we used the official open-source implementations provided by the authors. To ensure a fair comparison, we conducted limited grid searches around the recommended optimal hyperparameters in each model to adapt them to the characteristics of our fMRI datasets.

All models were trained under the same hardware environment, and early stopping was applied to prevent overfitting. For SPDNet, we adopted the SPDNet-2BiRe configuration, which contains two BiMap/ReEig blocks. The learning rate was fixed at $10^{-2}$, the batch size was set to 64, weights were initialized as random semi-orthogonal matrices, and the rectification threshold was set to $\varepsilon = 10^{-4}$. Training was performed for 100 epochs with early stopping. For the Brain Network Transformer, we used the Adam optimizer with a learning rate of $10^{-4}$ and a weight decay of $10^{-4}$. The batch size was set to 128, and training lasted for 100 epochs with early stopping. We followed the official implementation, using 4 attention heads, a feedforward dimension of 1024, ReLU activation, and dropout 0.1. For Mamba, we maintained the same backbone configuration as MAMBAxBrain across different tasks. The Adam optimizer (without weight decay) was used with a learning rate of $2 \times 10^{-4}$ and a batch size of 128. Each model was trained for 100 epochs with early stopping. For FST-Mamba, we used the AdamW optimizer with a learning rate of 0.001 and a batch size of 64. Training was performed for 100 epochs with a cosine annealing learning rate schedule and early stopping. For CorrNN, we used a simple baseline with one linear projection from the FC feature vector to a 256-dimensional hidden representation, followed by BatchNorm1d, and used the resulting features for downstream classification. To account for differences in temporal window length and sample size across tasks, we adjusted key hyperparameters according to the input sequence length, ensuring comparable model complexity across different tasks.

#### Impact of sampling length on model performance

Supplementary Fig. S1 illustrates the effect of resting-state (Rest) data length on the prediction accuracy across all eight tasks. Overall, as the Rest data sampling length increases from 400 to 4800, the prediction accuracy for all tasks shows an upward trend. The improvement in accuracy is most pronounced in the initial stage (lengths from 400 to 1000). Notably, the accuracy of the 'Rest' task itself is most sensitive to data length; it has the lowest accuracy at a length of 400 (approximately 74%) but rises rapidly with increasing data, reaching a top-tier level comparable to other tasks (>95%) at a length of 4800. When the sampling length exceeds 2400, the performance gains for most tasks tend to plateau, demonstrating a trend of diminishing marginal returns, which suggests that the model may have neared the saturation point for extracting effective information from the resting-state data.

#### Model efficiency and generalization of sparse connectivity features

By analyzing the weight distribution of the fully connected network, we identified 295 relatively important edges. Training the model with only these 295 edges led to just a 1.4% drop in performance compared to using all 6,441 edges, demonstrating that the model relies mainly on a sparse set of critical connections. Specifically, when trained on the HCP Young Adult dataset, the model achieved an accuracy of 93.213%. Furthermore, when the same trained model was evaluated on the HCP Test-Retest dataset, it still reached an accuracy of 81.536%, highlighting both its efficiency and cross-dataset generalization ability.

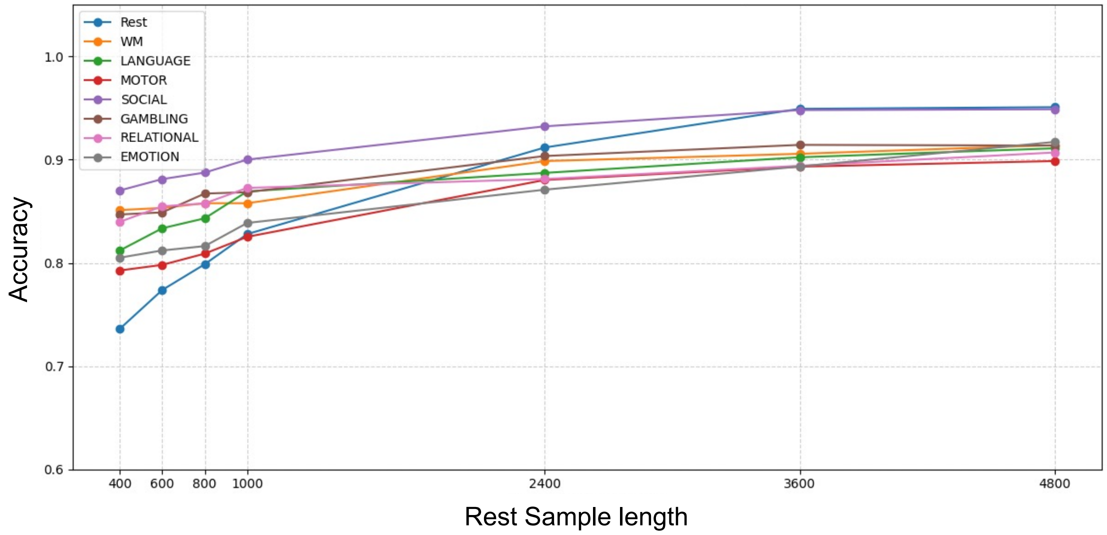


**Supplementary Figure S1 | Effect of resting-state data length on cognitive task decoding accuracy.**

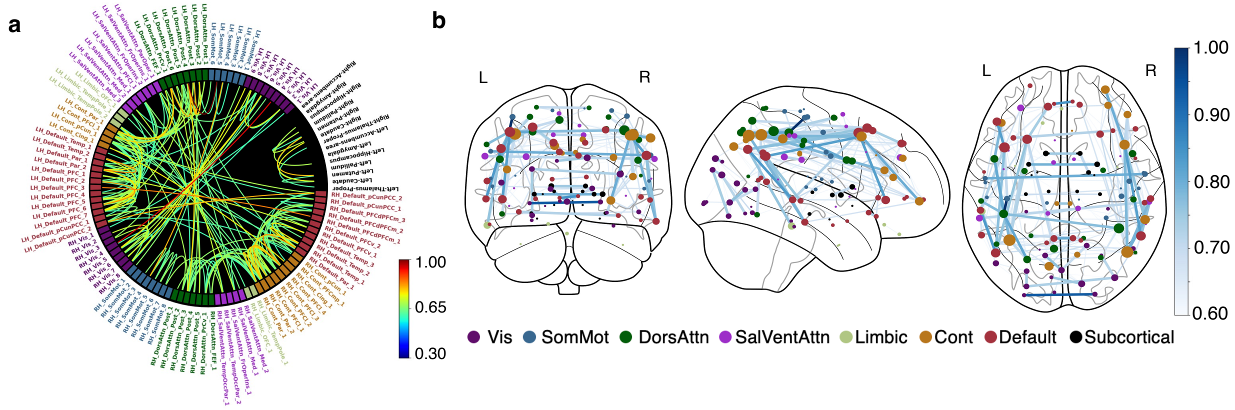


**Supplementary Figure S2 | Visualization of functional connectivity under joint training on all tasks.** **a** Connectivity circle representation illustrating inter-hemispheric and intra-hemispheric connectivity patterns (connection threshold set to 0.6). **b** Connectome plot projected onto real atlas coordinates, showing the spatial distribution of functional connections (threshold set to 0.6).

#### Robustness to sliding-window overlap

Supplementary Table S2 reports the stability of brain fingerprinting (FP) and cognitive task decoding (TD) to the sliding-window stride. Because a stride of 1 produces windows that overlap by 99 of their 100 TRs, we repeated both evaluations with non-overlapping windows (stride equal to the window length, 100 TRs, so that evaluation windows share no time points). Accuracy changes by less than one percentage point in every setting (the $\Delta$ row), both in-domain (HCP-YA) and across sessions (HCP-RT), indicating that the reported performance is not driven by the degree of overlap between adjacent windows.

Table S2: **Stability of fingerprinting (FP) and cognitive task decoding (TD) to the sliding-window stride.** Accuracy (%, mean $\pm$ standard deviation across repeated runs) under the default overlapping evaluation (stride 1) and a non-overlapping evaluation (stride equal to the window length, 100 TRs, so evaluation windows share no time points). The $\Delta$ row gives the change (non-overlapping minus overlapping) in percentage points; performance is essentially unchanged in every setting.

| **Evaluation protocol** | **In-domain (HCP-YA)** | | **Generalization (HCP-RT)** | |
|---|---|---|---|---|
| | **Brain FP** | **Cognitive TD** | **Brain FP** | **Cognitive TD** |
| Overlapping windows (stride 1) | 96.07±0.55 | 99.47±0.06 | 94.94±0.49 | 98.93±0.38 |
| Non-overlapping windows (stride = 100 TRs) | 95.42±0.88 | 99.31±0.12 | 94.31±0.74 | 98.67±0.41 |
| $\Delta$ (percentage points) | −0.65 | −0.16 | −0.63 | −0.26 |

#### Within-condition reaction time prediction

Reaction time (RT) prediction targets the condition-level mean RT of each HCP task condition. Because the six conditions differ systematically in mean RT (working memory, theory-of-mind, random, and relational conditions are comparatively fast, whereas math and story/language conditions are slow), a large part of the aggregate $R^2 = 0.92$ reflects the model separating conditions rather than individual neural variability. To quantify this, we computed $R^2$ separately within each condition. As shown in Supplementary Table S3, within individual conditions the model explains much less variance than the aggregate $R^2 = 0.92$ (mean within-condition $R^2 = 0.19$, range 0.12–0.27). Individual, within-condition RT prediction is therefore limited, which we report explicitly and note as a limitation.

#### Architecture vs. multi-task benefit at a matched data budget

Supplementary Table S4 reports, for every method, single-task training from scratch on the same 20% of each target task, together with few-shot adaptation of the multi-task MAMBAxBrain on that same 20% (see the main-text Results for interpretation). Among the single-task models MAMBAxBrain leads on every task, isolating the architectural advantage at the few-shot budget; few-shot adapting the multi-task model on the same 20% adds a further, larger gain (e.g., reaction time $R^2 \times 100$: 34.50 $\rightarrow$ 71.90), isolating the multi-task benefit at fixed architecture and data.

Table S3: **Within-condition reaction time prediction $R^2$.** $R^2$ computed separately within each HCP condition, compared with the overall $R^2$ across all conditions. The aggregate $R^2$ is dominated by between-condition differences in mean RT: within each condition the model explains substantially less variance.

| **Task** | $R^2$ |
|---|---|
| Story (Language) | 0.19 |
| Math (Language) | 0.23 |
| TOM (Social) | 0.12 |
| Random (Social) | 0.16 |
| WM | 0.14 |
| Relational | 0.27 |
| Mean within-condition | 0.19 |
| Overall (all conditions) | 0.92 |

Table S4: **Architecture vs. multi-task benefit at a matched 20% target-data budget.** Every method is trained single-task from scratch on the same 20% of each target task; among these single-task models MAMBAxBrain leads on every task, isolating the architectural advantage. The final row few-shot adapts the multi-task MAMBAxBrain on the same 20%, and its gain over single-task MAMBAxBrain isolates the multi-task benefit. FP and TD are accuracies (%); RT is $R^2 \times 100$.

| **Method (20% target data)** | **FP** (acc %) | **TD** (acc %) | **RT** ($R^2 \times 100$) |
|---|---|---|---|
| SPDNet | 33.50 | 79.20 | 15.80 |
| BrainNetTF | 39.70 | 85.30 | 21.60 |
| CorrNN | 58.40 | 88.10 | 25.90 |
| Mamba | 65.20 | 90.40 | 29.40 |
| FST-Mamba | 66.80 | 89.10 | 31.20 |
| **MAMBAxBrain (single-task)** | **70.92** | **90.30** | **34.50** |
| **MAMBAxBrain (multi-task few-shot)** | **84.20** | **96.50** | **71.90** |

#### Computational cost comparison

Supplementary Table S5 reports the computational cost of MAMBAxBrain and the baseline models. To make the comparison fair and to address the differing per-task state dimensions listed in Table 2 (which are MAMBAxBrain's own task configurations rather than a cross-model setting), all models were profiled under a matched input configuration (114 ROIs, window length 100) on the same hardware (a single NVIDIA RTX 3080 Ti), with MAMBAxBrain reported at its largest per-task state dimension (150) so that its cost is an upper bound, and timing measured for a batch of 128 samples. MAMBAxBrain reaches its accuracy at a cost close to the single-stream Mamba backbone (10.9 vs. 10.6 GFLOPs; 6.6 vs. 6.0 ms inference) and at roughly half the cost of the hierarchical FST-Mamba (10.9 vs. 28.0 GFLOPs; 26.3 vs. 60.0 ms per training step; 0.7 vs. 1.4 GB training memory). This confirms that the dual-stream design adds the functional-connectivity pathway without the heavy overhead of stacked state-space encoders, substantiating the efficiency statements made in the main text.

Table S5: **Computational cost across models.** Parameter count, multiply–accumulate operations (MACs), floating-point operations (FLOPs), inference time and peak memory, and training time per step and peak memory. All models were profiled under matched inputs (114 ROIs, window length 100) on a single NVIDIA RTX 3080 Ti, with MAMBAxBrain at its largest per-task state dimension (150); timing is reported for a batch of 128 samples.

| **Model** | **Params (M)** | **MACs (G)** | **FLOPs (G)** | **Infer** (ms/batch) | **Infer mem** (GB) | **Train** (ms/step) | **Train mem** (GB) |
|---|---|---|---|---|---|---|---|
| SPDNet | 0.4 | 0.6 | 1.2 | 80.0 | 0.2 | 300.0 | 0.3 |
| BrainNetTF | 1.6 | 8.8 | 17.6 | 8.0 | 0.4 | 32.0 | 0.6 |
| CorrNN | 0.9 | 0.2 | 0.4 | 1.5 | 0.1 | 5.0 | 0.2 |
| Mamba | 1.6 | 5.3 | 10.6 | 6.0 | 0.4 | 24.0 | 0.6 |
| FST-Mamba | 3.0 | 14.0 | 28.0 | 15.0 | 0.9 | 60.0 | 1.4 |
| **MAMBAxBrain** | 2.8 | 5.5 | 10.9 | 6.6 | 0.4 | 26.3 | 0.7 |

#### Subject-level stratified cross-validation for schizophrenia classification

To address the small size of the UCSF schizophrenia cohort and the possibility of within-subject leakage, we re-evaluated all models with diagnosis-stratified, subject-level 5-fold cross-validation, in which all windows of a given subject are confined to a single fold so that no subject appears in both training and test. Supplementary Table S6 reports accuracy and AUC (mean $\pm$ s.d. across folds). All models score lower than under the standard split and the fold-to-fold variability is large, consistent with the limited sample size; nonetheless MAMBAxBrain remains the best on both accuracy (0.82) and AUC (0.83), ahead of the strongest baseline (FST-Mamba, 0.76/0.80), indicating that its advantage on this task is not an artifact of a single favorable split.

Table S6: **Schizophrenia classification under diagnosis-stratified, subject-disjoint 5-fold cross-validation (UCSF).** Accuracy and AUC (mean ± s.d. across the five folds), with all windows of a subject confined to a single fold so that no subject appears in both training and test.

| Model | Acc | AUC |
|---|---|---|
| SPDNet | 0.68±0.12 | 0.70±0.19 |
| BrainNetTF | 0.74±0.10 | 0.78±0.16 |
| CorrNN | 0.75±0.10 | 0.79±0.15 |
| Mamba | 0.71±0.11 | 0.74±0.18 |
| FST-Mamba | 0.76±0.09 | 0.80±0.14 |
| **MAMBAxBrain** | **0.82±0.11** | **0.83±0.17** |

#### External validation on the COBRE schizophrenia cohort

To test whether the schizophrenia classification performance is specific to the UCSF cohort, we evaluated all models on the independent, publicly available COBRE dataset (72 patients with schizophrenia and 74 healthy controls). Models trained on the UCSF cohort were applied to the entire COBRE cohort at the subject level without any retraining, and no COBRE data were used for training. As shown in Supplementary Table S7, MAMBAxBrain achieves the highest external performance (AUC = 0.81, accuracy = 0.68), exceeding every baseline and indicating that the learned representations transfer to an independent site rather than reflecting dataset-specific artifacts. Consistent with the substantial cross-site domain shift (scanner, acquisition, and clinical sampling differences), absolute performance is lower than the within-cohort UCSF result, and sensitivity in particular drops to 0.43; the external setting is therefore best interpreted as evidence of moderate, above-chance transfer rather than of clinical-grade accuracy.

Table S7: **External validation on the COBRE cohort (subject-level).** All models were trained on the UCSF cohort and evaluated on the full, independent COBRE cohort (72 patients, 74 controls) without any COBRE fine-tuning. AUC, accuracy (Acc), sensitivity (Sens), and specificity (Spec) are reported.

| Model | AUC | Acc | Sens | Spec |
|---|---|---|---|---|
| SPDNet | 0.61 | 0.55 | 0.08 | 1.00 |
| BrainNetTF | 0.75 | 0.56 | 0.13 | 0.98 |
| CorrNN | 0.76 | 0.55 | 0.10 | 0.99 |
| Mamba | 0.76 | 0.53 | 0.06 | 0.99 |
| FST-Mamba | 0.78 | 0.60 | 0.26 | 0.93 |
| **MAMBAxBrain** | **0.81** | **0.68** | **0.43** | 0.92 |

#### Controlled component ablation

To isolate the contribution of each component, Supplementary Table S8 reports a controlled, change-one-thing-at-a-time ablation in which every variant keeps the rest of the architecture and the training budget fixed. Removing the FC stream (temporal stream only, which coincides with the Mamba backbone) and removing the temporal stream (FC stream only, which coincides with CorrNN) both reduce performance, most strongly for cross-session fingerprinting (94.94 $\rightarrow$ 83.26 and 86.12, respectively), confirming that the two streams are complementary. Replacing the learnable fusion with simple concatenation causes a smaller but consistent drop (e.g., in-domain fingerprinting 96.07 $\rightarrow$ 95.40), showing that the adaptive fusion contributes beyond plain feature stacking. Finally, replacing the Mamba temporal encoder with Transformer, TCN, LSTM, or GRU encoders of matched capacity degrades performance monotonically, with the largest gaps again under cross-session evaluation (cross-session fingerprinting 94.94 for Mamba vs. 91.10/90.20/88.50/87.90 for Transformer/TCN/LSTM/GRU). Together these results demonstrate that the selective state-space encoder, the dual-stream design, and the learnable fusion each contribute, and that the gains are not attributable to model capacity or the dual-stream scaffolding alone.

Table S8: **Controlled component ablation.** Each variant changes a single component while holding the rest of the architecture and the training budget fixed. In-domain (HCP-YA) metrics are schizophrenia accuracy (SZ), brain fingerprinting accuracy (FP), cognitive task decoding accuracy (TD), and reaction time ($R^2 \times 100$); cross-session (HCP-RT) metrics are FP and TD accuracy. "TS stream only" and "FC stream only" coincide with the Mamba and CorrNN baselines in Table 1. Values are mean ± standard deviation across repeated runs.

| **Variant** | **In-domain (HCP-YA)** | | | | **Cross-session (HCP-RT)** | |
|---|---|---|---|---|---|---|
| | **SZ** | **FP** | **TD** | **RT** ($R^2 \times 100$) | **FP** | **TD** |
| **MAMBAxBrain (full)** | **91.05±0.71** | **96.07±0.55** | **99.47±0.06** | **91.80±0.40** | **94.94±0.49** | **98.93±0.38** |
| TS stream only (≈Mamba) | 82.36±0.63 | 94.95±0.50 | 99.43±0.12 | 88.10±0.71 | 83.26±0.82 | 97.41±0.34 |
| FC stream only (≈CorrNN) | 86.78±0.99 | 93.80±0.13 | 98.99±0.09 | 88.09±0.65 | 86.12±1.78 | 97.05±0.91 |
| Simple concatenation | 90.10±0.85 | 95.40±0.60 | 99.38±0.12 | 90.50±0.55 | 93.30±0.75 | 98.65±0.45 |
| Mamba → Transformer | 89.40±0.90 | 95.00±0.65 | 99.28±0.15 | 89.50±0.70 | 91.10±0.95 | 98.45±0.50 |
| Mamba → TCN | 88.70±0.95 | 94.55±0.70 | 99.18±0.16 | 88.80±0.80 | 90.20±1.05 | 98.25±0.55 |
| Mamba → LSTM | 87.80±1.00 | 94.30±0.78 | 99.05±0.18 | 88.30±0.90 | 88.50±1.15 | 97.85±0.55 |
| Mamba → GRU | 87.40±1.05 | 94.10±0.82 | 98.98±0.20 | 88.00±0.95 | 87.90±1.20 | 97.75±0.58 |
